# Modelling framework to demonstrate elimination of a vector population: tsetse elimination in Chad

**DOI:** 10.1101/2025.09.09.675028

**Authors:** John Hargrove, Mahamat Hissene Mahamat, Moukhtar Aldjibert, Wilfrid Yoni, Djoukzoumka Signaboubo, Justin Darnas, Ernest Salou, Inaki Tirados, Albert Mugenyi, Priscille Barreaux, Philippe Solano, Antoine Marc Gaby Barreaux

## Abstract

Every year, over 700 000 people, particularly children under five, die from vector-borne diseases worldwide. Effectively controlling current endemics and preventing new outbreaks requires an integrated approach that can lead to the elimination of both vectors and diseases. In the last two decades, integrating medical interventions and vector control has significantly reduced the incidence of Gambian Human African Trypanosomiasis (gHAT), with the World Health Organization validating eight countries as having eliminated the disease as a public health problem. However, elimination of the tsetse vector has not been confirmed, leaving the possibility of re-emergence. We developed a five-step modelling framework to assess vector elimination by calculating: (i) the probability of vector capture; (ii) the probability of observing a series of zero catches, even without actual elimination; (iii) the probability of natural elimination; (iv) the probability of failing to detect a rebound; and (v) the reinvasion risk. Our case study is g-HAT in Mandoul, Chad and the elimination of *G. fuscipes fuscipes*. We used vector control from 2014 to 2025 with no tsetse detected since 2018. We cannot yet conclude, with more than 90% confidence, that tsetse has been eliminated from Mandoul, nor that any remnant population will be naturally eliminated. However, since vector control was stopped in April 2025, we estimate that with continued sampling over the next two years, and no tsetse detected, elimination could be demonstrated with 99% confidence. Our multi-step modelling framework can be applied to other vectors, providing policymakers with clear guidelines for ongoing and future efforts.

**Significance Statement:** The World Health Organisation has set the elimination of transmission of several neglected tropical vector-borne diseases, including human African trypanosomiasis (sleeping sickness), as a target for 2030. We show that deliberate elimination of tsetse, the vector, is feasible and can be demonstrated. We draw on our large-scale intervention in Mandoul, Chad where 3000 insecticide-treated Tiny Targets were deployed between 2014 and 2025, with no tsetse detected since 2018. While small undetected remnant populations cannot be entirely excluded, they would rapidly rebound in the absence of control, rendering them detectable. If no tsetse are caught over the next two years, it will confirm elimination. This illustrates a pathway for assessing and achieving vector elimination as a cornerstone of disease eradication.

## Introduction

Vector-borne diseases (VBDs) remain a major public health challenge worldwide with over 700 000 deaths annually, particularly among children under five (1). Sub-Saharan Africa bears the heaviest burden, where malaria, dengue and other neglected tropical diseases (NTDs) affect millions of women and men. To successfully interrupt disease transmission, control approaches must address critical questions: when, where, how and with whom to act. When implemented effectively, integrated strategies can eliminate diseases to the point where the World Health Organization (WHO) validates them as Eliminated as a Public Health Problem (EPHP) (2).

Recent progress includes EPHP validation for gambiense Human African Trypanosomiasis (g-HAT or sleeping sickness), in Chad in 2024 (3), alongside seven other endemic countries including Guinea (4), and Côte d’Ivoire (5). After EPHP, the WHO goal is elimination of g-HAT transmission by 2030, with the aim of achieving zero cases of the disease by that time (6). Integrated approaches must target not only diagnostics, vaccines and therapeutics for the parasite and host, but also target arthropod vectors to reduce host-vector contact, and ultimately reduce and even eliminate transmission (5, 7–10).

Vector control successes include insecticide treated bednets and innovative housing approaches to control mosquitoes (7, 11) and Tiny Targets to control tsetse (*Glossina spp*), the sole vectors of HAT (12). In various settings and countries, these small insecticide impregnated pieces of fabric, Tiny Targets, have significantly reduced tsetse populations, lowering human-vector contact (5, 13, 14) and helping to halt g-HAT transmission (15). Vector ecology also strongly influences VBD epidemiology (16) and vector populations are highly sensitive to environmental and global change factors (17–21) which affect vector elimination prospects.

Despite progress, a critical question remains: how to know when elimination has been achieved and how to quantify the risk of error. While disease cases or vector captures may reach zero, confirming the complete elimination of either is a far more complex issue. The challenge lies in determining whether the absence of disease or vectors truly reflects elimination, or simply a limitation in our surveillance and detection capabilities. In some cases, the cessation of control efforts has led to a rebound in disease transmission. In some instances, stopping control has led to resurgence: halting control in the 1960s caused an estimated 300,000 cases of g-HAT in the 1990s, underscoring the need for a more robust understanding of vector population dynamics (2).

To address these challenges, we propose a novel mathematical and modelling framework, addressing the three core questions of VBD elimination: (i) Can elimination, in theory, be achieved? (ii) How will it be attempted in practice? (iii) How do we know when it has been achieved?

In our study, using tsetse and HAT as an example, we focus on the third question, considering diseases and vectors that can be eliminated with currently available methods (diagnostics, treatments and tiny targets for vector control) and that for tsetse elimination feasibility can be estimated from birth and death rates and the size of the starting population (22–24).

When a VBD elimination program reaches a stage where zero cases of disease are diagnosed, and/or zero vectors are captured, elimination of the disease and/or the vector may indeed have been achieved. Alternatively, this outcome may simply reflect the lower detection threshold of our sampling methods, below which parasites or vectors remain undetectable. If control efforts are halted while the disease and/or vectors persist – though undetectable with current tools – there is a risk of renewed transmission, rising case numbers, and ultimately a re-emergence of the vector, and/or the disease. We therefore investigate the theory underlying mathematical assessment of the probability that elimination has truly been achieved. We primarily focus is on vector elimination, though clear parallels exist for disease elimination. Specifically, we explore mathematical approaches to estimate the risk incurred when declaring a vector population eliminated. This can be structured as a five-steps approach and modelling framework:

I. Before vector control is started, estimate the probability of capturing a vector with the surveillance tools to be used. This may be informed by existing literature when available, or by baseline mark-recapture, or other studies, to calculate the probability of capturing a vector. Seber (25) provides an excellent review of available methods and see Williams et al. (26) for more recent advances in the field.
II. Calculate the conditional probability of observing a series of zero catches given that at least one individual vector is still present. If that probability is sufficiently low, we conclude the vector has been eliminated (23);
III. If the vector population is very low but not yet eliminated, calculate the probability of natural extinction, purely by chance, without further control efforts (22, 24);
IV. Use growth models to estimate the expected vector population at various times after the cessation of control efforts, assuming survival of at least one reproductive female. Failure to detect a rebound in the vector population after extended periods supports elimination (23).
V. Calculate the probability of vector reinvasion to further support the demonstration of vector elimination (27).

As a case study of vector elimination modelling, we examine tsetse-transmitted g-HAT focusing on efforts to eliminate the tsetse species *Glossina fuscipes fuscipes* from the Mandoul focus in southern Chad (13) – a site of serious g-HAT outbreaks. Both sexes of the tsetse fly are strictly hematophagous and are thus both capable of transmission. The disease is of particular interest because, while massive outbreaks occurred over the last two centuries, g-HAT seems now to be under control in many countries, with EPHP validated (3–5) and even elimination of transmission in some cases.

Tsetse provide a relatively simple, and tractable system for estimating the probability of vector elimination because of their slow, and predictable reproductive biology. Each adult female produces, only one larva every 9-11 days (28, 29). The larva nearly the same weight as its postpartum mother, contains the energy, microbiome and materials required to pass through all of the final larval and pupal, stages of metamorphosis, into an adult-sized teneral fly (28, 30, 31). Following larviposition, the free-living pupa does not feed at all. Reproduction continues year-round, independent of the availability of environmental water (32). The production of a single offspring at each birth event, separated by roughly 10 days, means that the fate of individual offspring may be regarded as independent, which is an important assumption for probabilistic modelling. The low reproductive rate also means that population growth rates become negative if adult female mortality exceeds 4% per day. Sustained mortality at or above this level leads to elimination (33). In principle, tsetse populations should therefore be relatively easy to eliminate, particularly since they have never shown resistance to insecticide and their simple life cycle makes it feasible to calculate the probability that elimination can indeed be achieved.

Our research provides a modelling framework for assessing the likelihood of vector elimination, using *G. f. fuscipes* and g-HAT in Chad as a case study. This work contributes to ongoing elimination strategies by evaluating whether current evidence is sufficient to demonstrate that specific vectors, such as tsetse, have been successfully eliminated.

## Results

### Step 1: probability of capturing one G. f. fuscipes in a biconical trap in Mandoul

Using data from Big Chamaunga Island, Lake Victoria, Kenya, we estimated the daily probability *p* that a female *G. f. fuscipes* in Mandoul is killed by a target or captured by an individual trap.

On Chamaunga, catches of female *G. f. fuscipes* declined 10^4^-fold in 1 year (Figure 1) after deployment of 30 Tiny Targets along 1.5 km of shoreline habitat (14). Following Hargrove (33), this rate of decline is consistent with targets killing a proportion *p* = 0.04 (*i*.*e*., 4%) of adult females per day. Accordingly, the proportion *q* = 1-*p* that is not killed each day by any of the 30 targets is 0.96. The probability that a single target fails to kill a single fly in a day is then 0.96^30 = 0.99864 and the proportion of the total population killed by one target is 1 - 0.99864 = 0.0014 or 0.14% per day. As a single biconical trap is estimated to catch about half as many tsetse as a Tiny Target (Esterhuizen et al., 2011), each trap should then capture (0.14/2) = 0.07% of the total fly population on the island per day.

**Figure 1.**
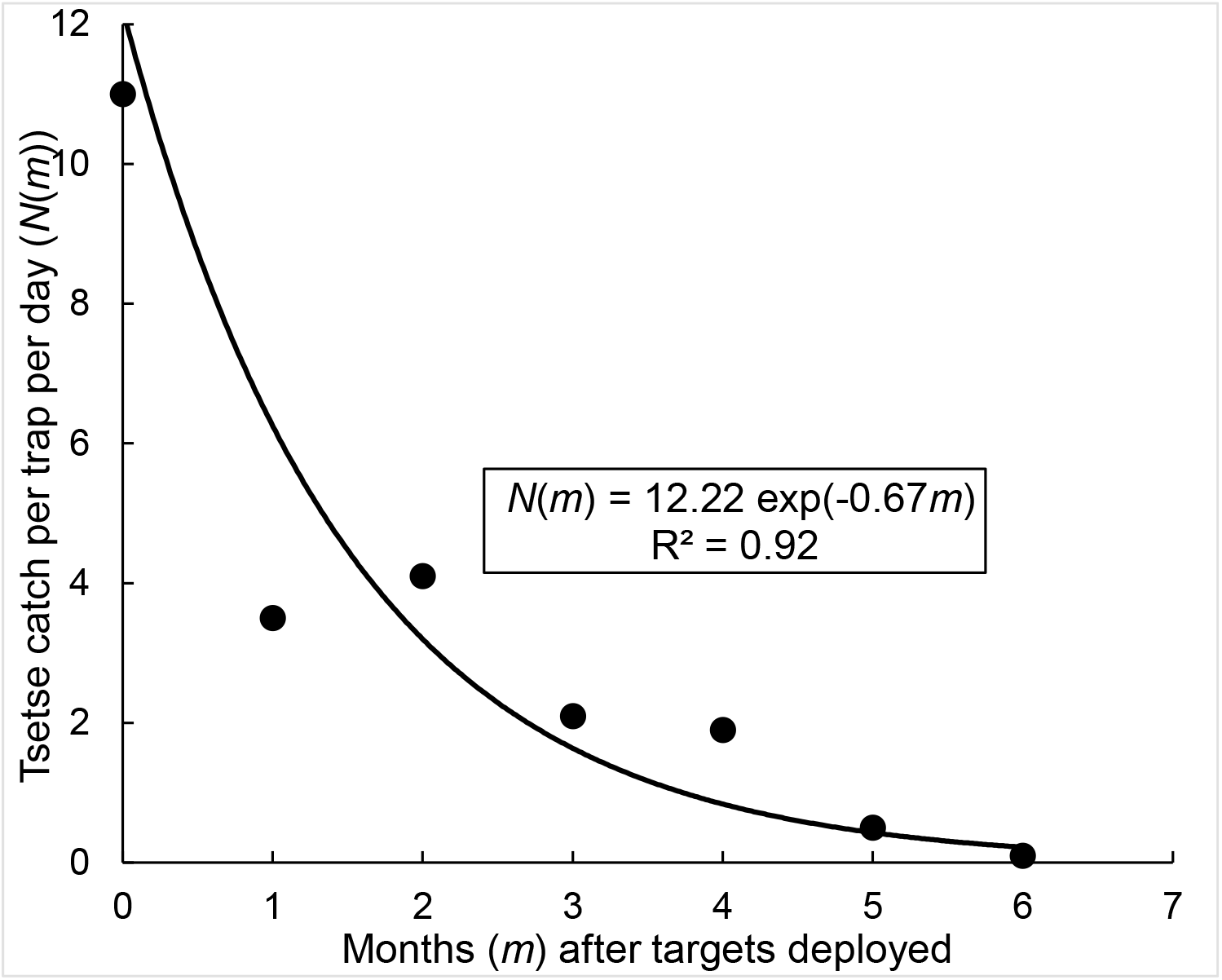
Decline in numbers of female G. f. fuscipes trapped on Big Chamaunga Island, following the deployment of 30 Tiny Targets on the 1.5 km perimeter of the island.

We assume these kill and recapture rates apply generally to riverine tsetse when using Tiny targets and such biconical traps, including the Mandoul populations, independent of habitat area. Implicit in this assumption is that targets cover all available habitat, such that each target’s 0.14% contribution is drawn from a local fraction of the total population. By the same logic, zero catches from a given trap reflect only the absence of tsetse in the immediate neighborhood of that particular trap.

In Mandoul, 145 *G. f. fuscipes* were captured in the 2013 baseline survey, and following the initial deployment of 2713 Tiny Targets in March 2014, the follow up surveillance sessions –in April and June 2014 – resulted in only 2 and 3 flies being captured respectively (13) – suggesting that the population had already declined by 98% (Figure 2). The decline was so rapid that it is difficult to estimate the true rate of decline of the population (Figure 3). Nonetheless, the results are consistent with a population growth rate of −0.767 per month, equivalent to a yearly decline of exp(−0.767^12^) ≈ 10^-4^ per year, and the overall control effort killing c. 4% per day of the adult female population – as for the Big Chamaunga Island. Hence each target in Mandoul killed (1 - 0.96)^(1/2713) = 0.0015% flies per day and if the biconical traps used to sample *G. f. fuscipes* is about half as efficient as the Tiny Targets (12), each trap is expected to catch about 0.00075% per day of all flies in the Mandoul control area. The probability of capturing one *G. f. fuscipes* in a biconical trap in Mandoul is of 0.0000075.

**Figure 2.**
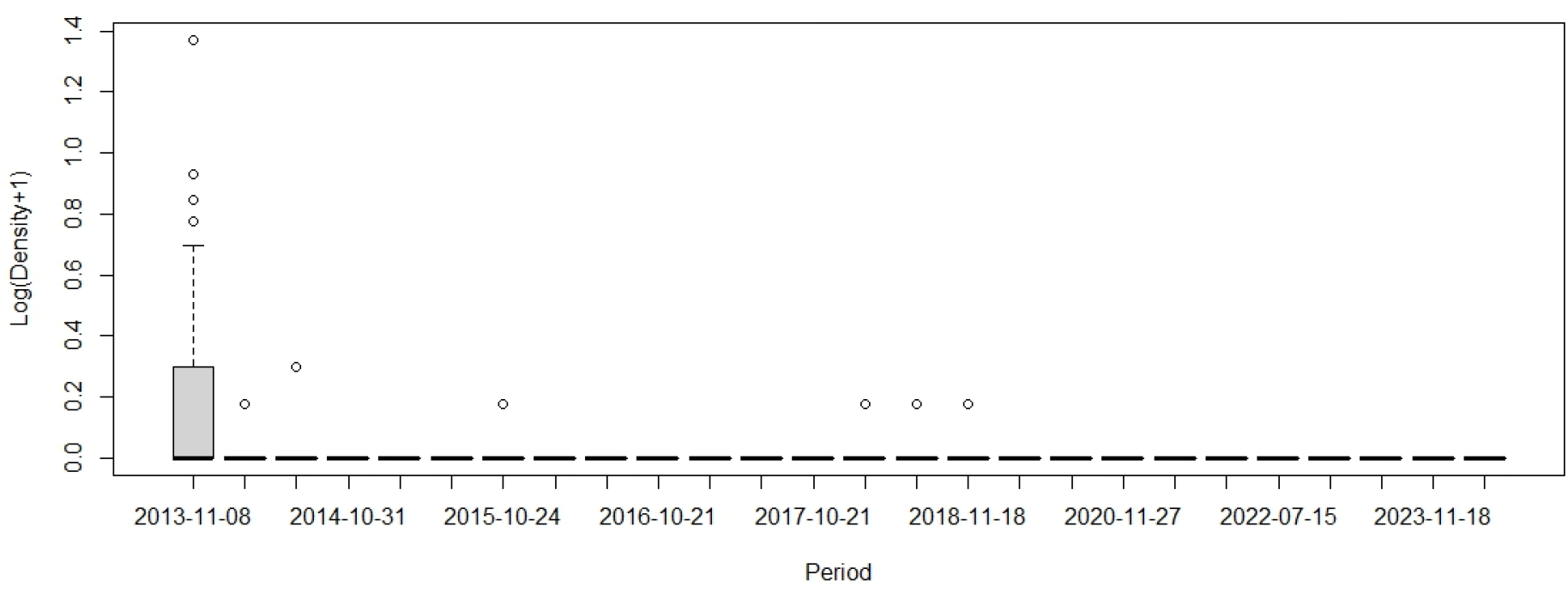
Catches of G. f. fuscipes from traps deployed, between November 2013 and October 2023, in the Mandoul focus of southern Chad. Numbers are plotted as the log (base 10) of the catch; zero catches cannot thus be plotted.

**Figure 3.**
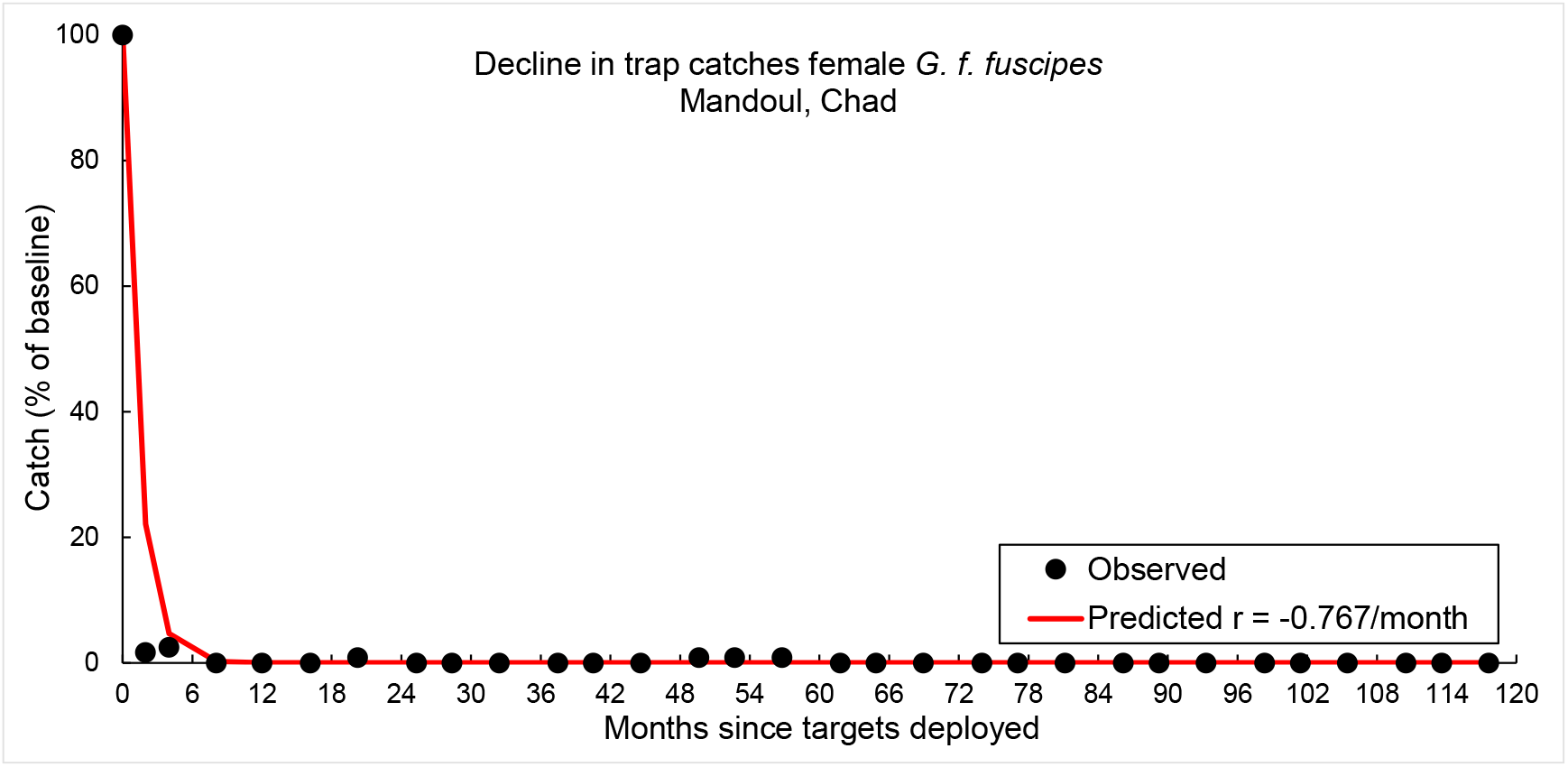
Observed catches of adult G. f. fuscipes in biconical traps in Mandoul, and the expected numbers if the growth rate was taken to be −0.767 per month.

Since this level of mortality in Mandoul should guarantee eventual elimination, we expect that the use of high densities of Tiny Targets tsetse should lead to the elimination of the Mandoul *G. f. fuscipes* population. The following sections assess the probability that elimination has indeed been achieved – or, conversely, the risk that elimination has not been achieved, despite continued failure to detect any flies using the above calculated probability of capture per trap.

### Step 2: Probability of zero tsetse catches, as a function of the numbers of tsetse surviving

Suppose now that there was only a single tsetse remaining in Mandoul, and that the area is sampled using 44 biconical traps, every day for 30 days. The probability of observing zero catches from every trap on every day is then ((1-0.0000075)^44^)^30^ ≈ 0.991, so a 99.1% chance of having a false absence/failing to catch that surviving fly. Similar calculations show that even if the 44 traps were run for 160 days, there would still be a 90% chance of failing to catch the surviving fly. Moreover, even with a 4-fold increase in trap efficiency, there would still be a 90% chance that 80 days of trapping would fail to catch a surviving fly (Figure 4A).

**Figure 4.**
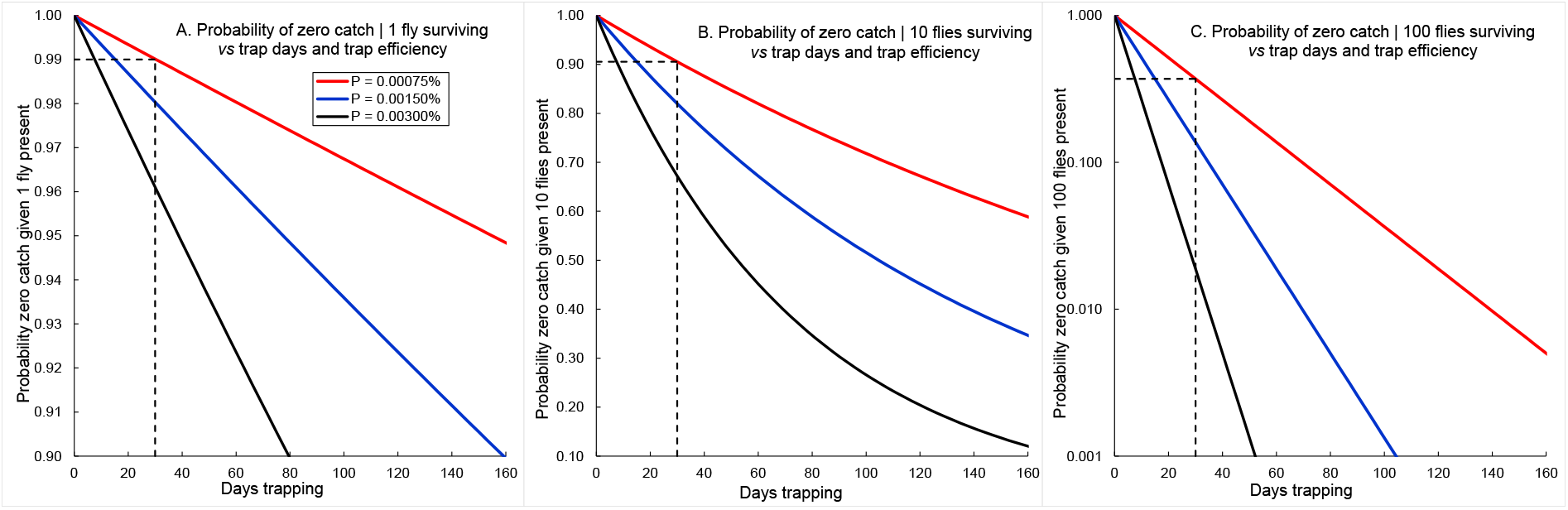
Probability of zero catch of tsetse, given that 1, 10 or 100 tsetse (A, B and C from left to right) have survived in the Mandoul area, as functions of trap efficiency and numbers of days trapping. The dotted lines show examples where 30 traps, each with 0.00075% efficacy, are run for 44 days.

If the number of tsetse surviving were 10, or 100, then the probability of observing a series of zero catches when some flies are still present obviously diminishes (Figure 4 B, C). With 10 surviving tsetse, 44 traps run for 30 days would still have a 90% chance of returning a series of zero catches/failing to catch any tsetse (Figure 4B): and, with 100 surviving flies, there would still be a 37% chance of failing to capture any.

### Step 2’: pursuing vector control and vector surveillance until one is 99% confident of tsetse elimination by relying on probability of capture in surveillance tools

If one flies remain and vector control is continued and vector monitoring still relies on the same 44 biconical traps, then it would take 13,900 days to be 99% confident that one has not missed capturing that single surviving fly. Even with 4 times more efficient traps it would still take 3475 days.

If 10 flies remain it would still take 1400 days with our current trap, and 348 days with a four times more efficient trap for a 99% confidence in tsetse elimination. Finally, if 100 flies remain, it would take 140 days with our current trap and 35 days with a four times more efficient trap to be 99% confident of tsetse elimination.

### Step 3: Probability of natural elimination of a tsetse remnant population in Mandoul

The previous sections underline the risk of inappropriately interpreting even a long series of zero trap catches as proof that the *G. f. fuscipes* population in Mandoul has been eliminated. If, however, we can at least be confident that the surviving population is small (10 or less) then what are the chances that such a population will be eliminated by chance? The results in Figure 5 suggest that, in general, tsetse populations are remarkably resilient to being eliminated by chance.

**Figure 5.**
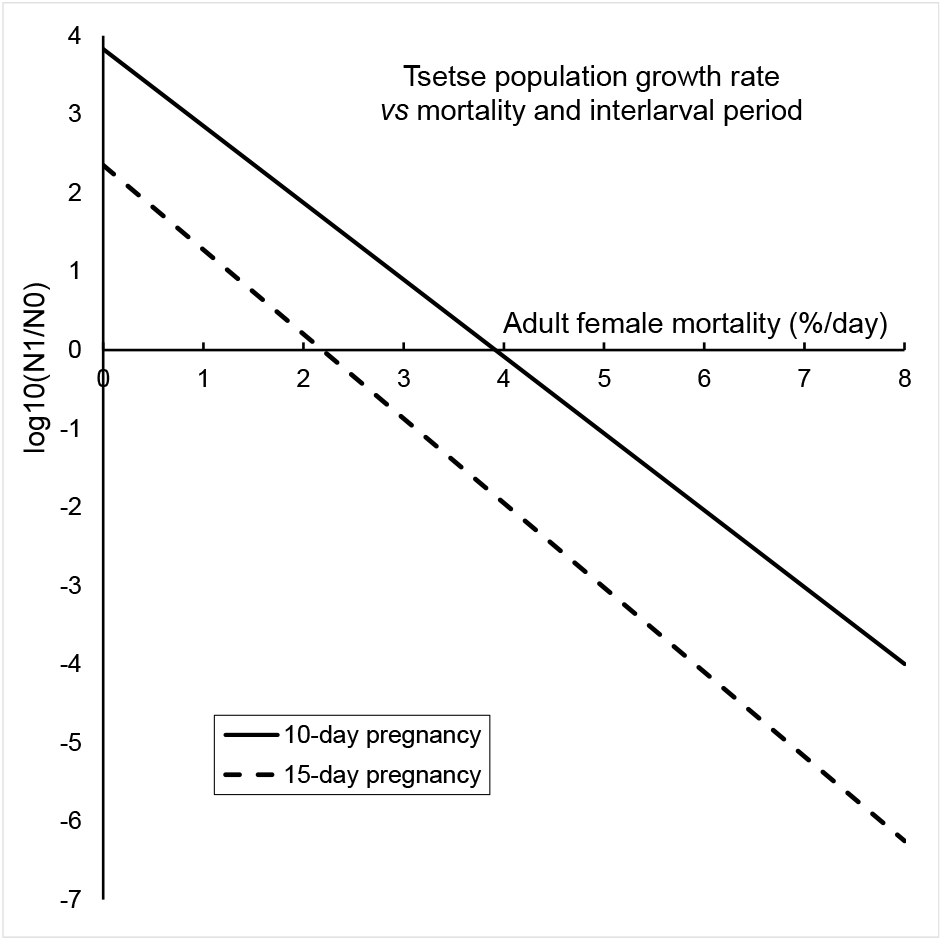
Annual growth rates of a tsetse population as a function of adult mortality and fecundity. Redrawn from “Hargrove J.W. (1988) Tsetse: the limits to population growth. Medical and Veterinary Entomology, 2, 203-217.”.

For the Mandoul population, if indeed adult female mortality could be maintained at 4% per day, then any population would be eliminated. If, however, the imposed mortality fell to, say 2% per day then the probability that a remnant population of 10 inseminated female tsetse disappears by chance is <10%. And even a remnant of 16 such flies is only eliminated by chance with probability <10% if female adult mortality is about 3% per day. In general, therefore, we would be unwise to rely on a remnant population being eliminated by chance in Mandoul.

### Step 4: Allowing for the detection of a post-vector-control rebound

The expected growth rate (i.e. rebound) of the hypothetical remaining tsetse population, following the removal of the Tiny Targets, can be inferred from Figure 5. At a low starting population number, implied by the inability to catch any tsetse with the surveillance system in place, we may safely assume that density-dependent mortalities among pupae and adults will be at a minimum. If we suppose that female mortality is at a conservative 2% per day, and that there are negligible losses among pupae, then Figure 5 suggests that the population could increase by up to 100-fold in the first year. Such rates of increase have been approached for field populations of *G. pallidipes* in Kenya (34) and *G. m. morsitans* and *G. pallidipes* in Zimbabwe (35). We might then expect that the *G. f. fuscipes* population in Mandoul could grow to a minimum of 1000 flies in two years. Even if the population increased by only 10-fold per year, the population should reach the 1000 level in three years.

If the Mandoul population were to rebound to a level of 1000 flies, 44 traps run for 3 weeks (21 days) would catch at least 1 fly with 99.9% certainty (Figure 6). That is to say, the probability of catching zero tsetse would be 0.001, or 0.1%. A zero catch over the whole 3-week period would then imply there was negligible risk in concluding that elimination had been achieved. The above scenario – based on the assumption of sampling a population that was assumed to reach the level of 1000 flies – might take two to three years to provide a decision that elimination has been achieved.

**Figure 6.**
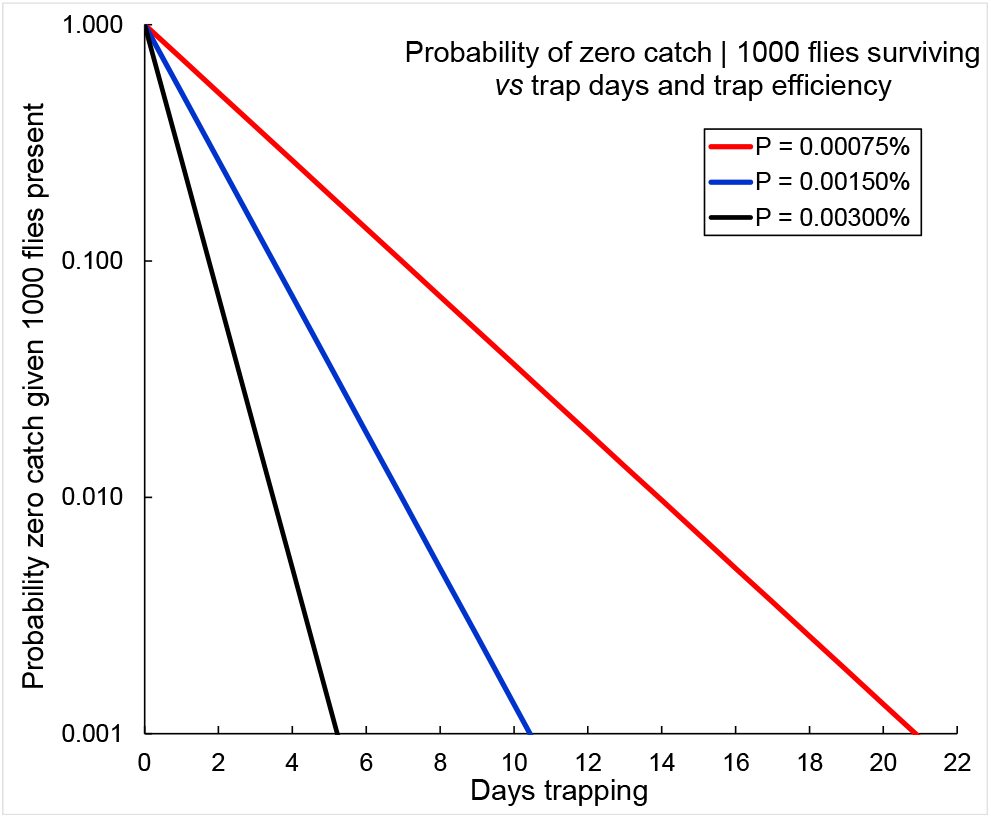
The probability of observing a series of zero catches of G. f. fuscipes in Mandoul after the removal of all Tiny Targets – given that the population has increased to a total of 1000 flies before the sampling effort is initiated – plotted as a function of trap efficiency and numbers of days of trapping.

The results in Figure 7 suggest that a demonstration of elimination could be achieved in much less than 3 years. If the sampling procedure continues to involve the use of 44 traps, the initial probability of catching a fly will be small – and one would have <90% confidence that elimination had been achieved elimination even after more than a year (550 days) of consecutive zero trap catches (Figure 7). Thereafter, however, confidence levels grow very rapidly – reaching 90% and 95% by days 600 and 650, respectively, and 99% after 2 years.

**Figure 7.**
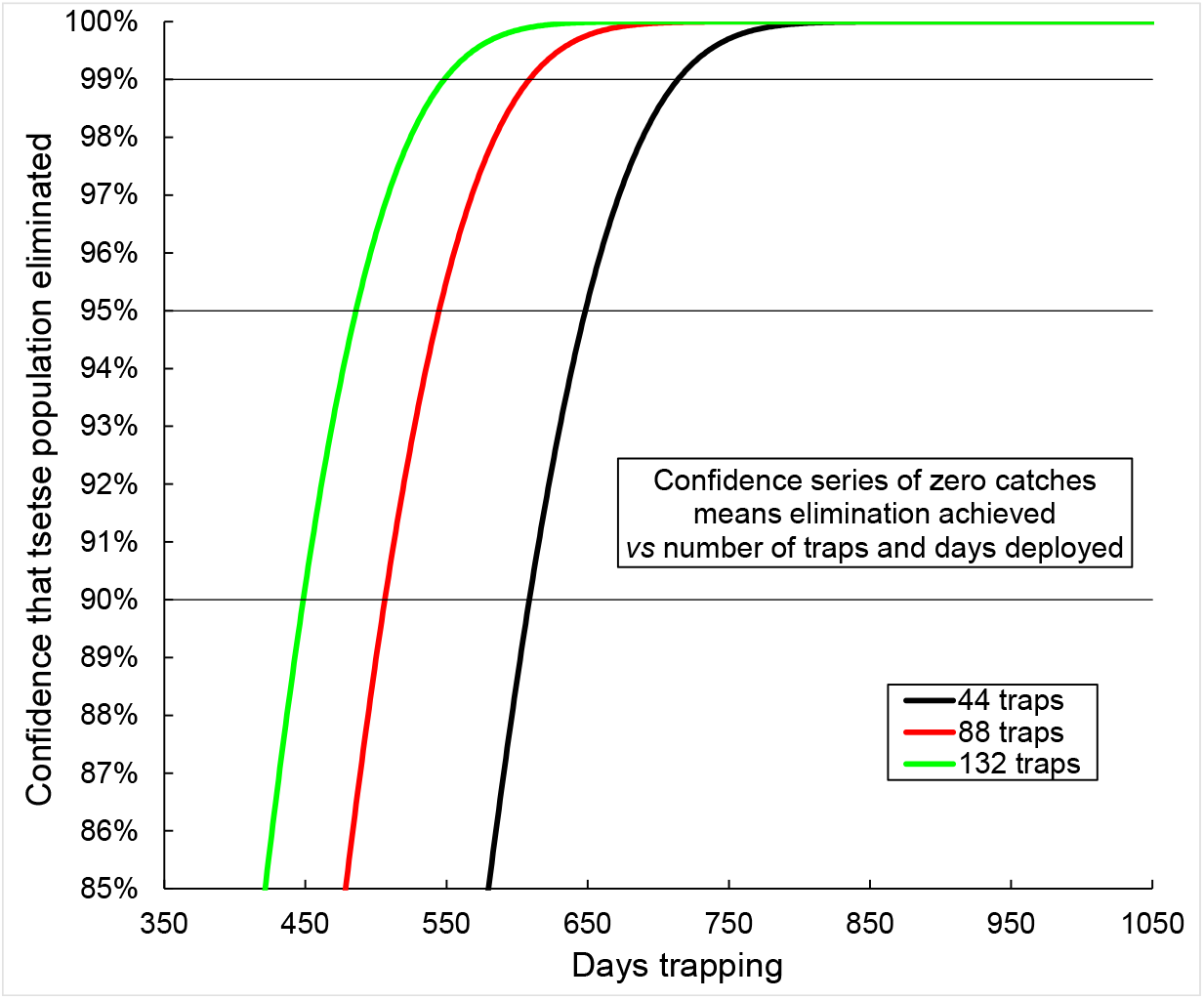
Estimates of confidence that series of zero trap catches support a conclusion that the Mandoul population of G. f. fuscipes has been eliminated. Calculated as a function of the number of traps used and the number of days sampling.

Doubling the sampling effort, by using 88 traps every day, makes only a modest difference to the outcome; 1.5 – 2 years of zero catches would still be required to be to be 99% confident that elimination had been achieved. Increasing the trapping effort further would probably be counterproductive. With 132 traps run daily, one could be 99% confident of elimination after 18 months of zero catches (Figure 7) but only if the traps were acting independently of each other. This assumption becomes increasingly unlikely as trap density increases (36).

### Step 5: Vector elimination and reinvasion risk

The closest population of tsetse, *G. f. fuscipes*, beyond the borders of Mandoul is in the neighborhood of Timbéri, 50 km distant. We estimate the probability that a tsetse fly could survive long enough to move between the two populations. We use the results of Hargrove & Lange (27) in modelling tsetse dispersal as diffusion in the plane. The fly’s position (*x, y, t*) in time and space is then defined by a normally distributed random variable with density function and if the population are widely separated, relative to the rate of diffusion, the equations can be much simplified.

For our study, Mandoul and Timbéri are separated by order 50 km, and the relative rate of diffusion is of the order of (0.04km)^2^/day, so that, for *t* <=100 days, (c-a)/(*kt*)^0.5^ ≈ 50/(0.04*t*)^0.5^ ≥ 10 >> 0. That being the case, we can make the required simplifications and the probability is then 0 (see equation 5 in material and methods). There is no chance that a fly can move between Timbéri and Mandoul in under 100 days, even if it survived for that period. For *t* > 100 days, the probability of completing the journey increases, but the probability that the fly survives this period decreases very rapidly. The danger of reinvasion due to diffusive movement is thus vanishingly low in any case.

## Discussion

No *G. f. fuscipes* have been captured in Mandoul since 2018, following a vector control program that began in 2014 and ended in 2025. It is not yet possible to conclude, with more than 90% certainty that the population has been eliminated, owing to the low probability of catching *G. f. fuscipes* with current trapping methods. Nonetheless, satellite images and community reports (source: Mahamat M.H., Aldjibert M., and Yoni W., 2025, pers. comm.) strongly suggest that tsetse no longer pose a threat: crops are cultivated in areas previously infested with tsetse and new human settlements are emerging along the Mandoul river. These developments further support the decision by the Chad tsetse control team to remove all Tiny Targets in early 2025. If current levels of surveillance continue for two more years without detecting a single *G. f. fuscipes* – then the authorities will be able to declare elimination that we will have demonstrated with 99% confidence. This represents one of the few modern examples of upcoming successful vector elimination in the world.

If a small number of tsetse persist, their population should increase once the mortality imposed by the Tiny Targets is lifted, making detection more likely. This represents a risk-reward situation: while risks appear low given over six years without captures, Tiny targets could be redeployed to quickly suppress any remnant population. The rewards, by contrast, are substantial – cost savings from halting control and the prospect of eventually ending all monitoring. Without stopping control, it would take an impractical 40 years of continuous trapping with current methods to reach 99% confident of elimination, and even longer with the present regime of a few trapping days per year. The near-zero risk of reinvasion from neighboring tsetse populations, combined with the improbability of accidental introduction via livestock or motorized transport, further supports cessation of control. No other tsetse population or species appear to be able to recolonize the area vacated by *G f. fuscipes*.

Our study highlights the difficulty of proving complete elimination of an isolated vector population when monitoring tools cannot reliably detect rare individuals. The problem scales with area size, as shown in the large-scale efforts to eliminate *G. m. centralis* in Botswana. There, contiguous blocks of 16,000 sq km in the Okavango Delta were sprayed in 2001 and 2002, and no tsetse have been caught since the completion of the second year of spraying (37). However, trap and fly-round sampling methods would not have detected tsetse at population densities of 1 per sq km, nor could such sampling methods cover the full habitat. The conclusion of elimination relied on the expectation that any surviving tsetse would have rebounded to detectable levels within two years.

Historically, such a “wait and see” approach relying on years without tsetse captures or reported cases of disease has underpinned declarations of elimination, with the passage of time without new cases providing sufficient evidence of local eradication. This is the basis for accepted cases such as *G. p. palpalis* on from Principe (38), *G. pallidipes* in Zululand (39) and *G. m. morsitans* in Umfurudzi Game Area of Zimbabwe (32) where elimination was inferred without long-term systematic sampling. The passage of years, and then decades, in which no tsetse and no cases of trypanosomiasis were reported, simply made it clear that the flies had indeed been eliminated. In contrast, very small or isolated areas allow quicker demonstration of elimination, as shown for *G. pallidipes* and *G. m. morsitans* on Antelope Island, Zimbabwe (35) and *G. austeni* on Unguja Island (40). Mathematical approaches, including landscape genetics (41) or species distribution modeling (42) have also been developed to help identify such target areas for disease or vector elimination.

Regardless of the scale or surveillance strategies, our framework provides a formal, probabilistic method to demonstrate vector elimination. This creates direct linkages to existing international processes: WHO’s elimination of public health problem (EPHP) (2) and elimination of transmission (EoT) for human African trypanosomiasis (HAT), FAO’s Progressive Control Pathways for Animal African Trypanosomiasis (AAT) (43), and WOAH’s recognition of AAT-free status. All are aiming for a greater integration of mathematical frameworks to move beyond reliance on the “wait and see” paradigm. Discussions are underway on incorporating our approach into official WHO and FAO procedures.

Although developed for tsetse, the framework is general and applicable to any vectors or even populations of beneficial species, be it plants or animals, at risk of extinction. Our results are therefore relevant to public health experts, policy makers, and conservation practitioners. In estimating the probability that the Mandoul population of *G. f. fuscipes* has been eliminated, we did not account for the possibility that long-term use of Tiny Targets may have induced behavioral change in tsetse, such as avoidance of these attractive systems. Similarly, selection pressure could have resulted in surviving flies feeding preferentially on host animals that can be accessed with reduced movement, thereby lowering the likelihood that flies are killed by Tiny Targets. There is, however, no evidence that suggest such behavioral or ecological shifts have occurred, and any potential impact on our results would be minimal. Importantly, the selection pressure imposed by our 44 surveillance traps is far weaker than that imposed by 2713 Tiny Targets, meaning that monitoring is less likely to be affected by the evolution of resistance than the vector control efforts.

We also adopted a conservative approach in our calculations to minimize the risk of incorrectly demonstrating elimination in Mandoul and to ensure our conclusions are not overly optimistic. For instance, in calculating the probability that a small surviving population of tsetse would not be eliminated by chance, we disregarded the possibility that adult virgin females might die before successfully mating, a factor that would further reduce the chance of persistence. In this respect, our estimates represent a worst-case scenario of the results obtained by the tsetse control team in Mandoul. This conservative approach strengthens our argument that the risks of removing the Tiny Targets are outweighed by the benefits of that policy.

## Conclusion

Whether or not tsetse have already been eliminated from the Mandoul area, Tiny Targets have successfully reduced tsetse population densities by several orders of magnitude to undetectable levels. In 2024, WHO validated elimination of gHAT as a public health problem in Chad, marking the operation a success (3). With the removal of Tiny Targets, continued monitoring will provide a definitive answer regarding the elimination of tsetse in the Mandoul region. More broadly, the multi-step theoretical approach developed in this study offers a rigorous method for demonstrating vector elimination. This framework is applicable to other disease vectors and provides guidance for policymakers and health authorities in planning, evaluating and sustaining future elimination efforts.

## Materials and Methods

### Modelling framework for assessing vector elimination

Using a decision tree (Figure 8), published methodology (22–24) and our novel modelling framework, we use up to five steps, in sequence, to estimate the probability that a vector population has been eliminated in a specific area:

**Figure 8.**
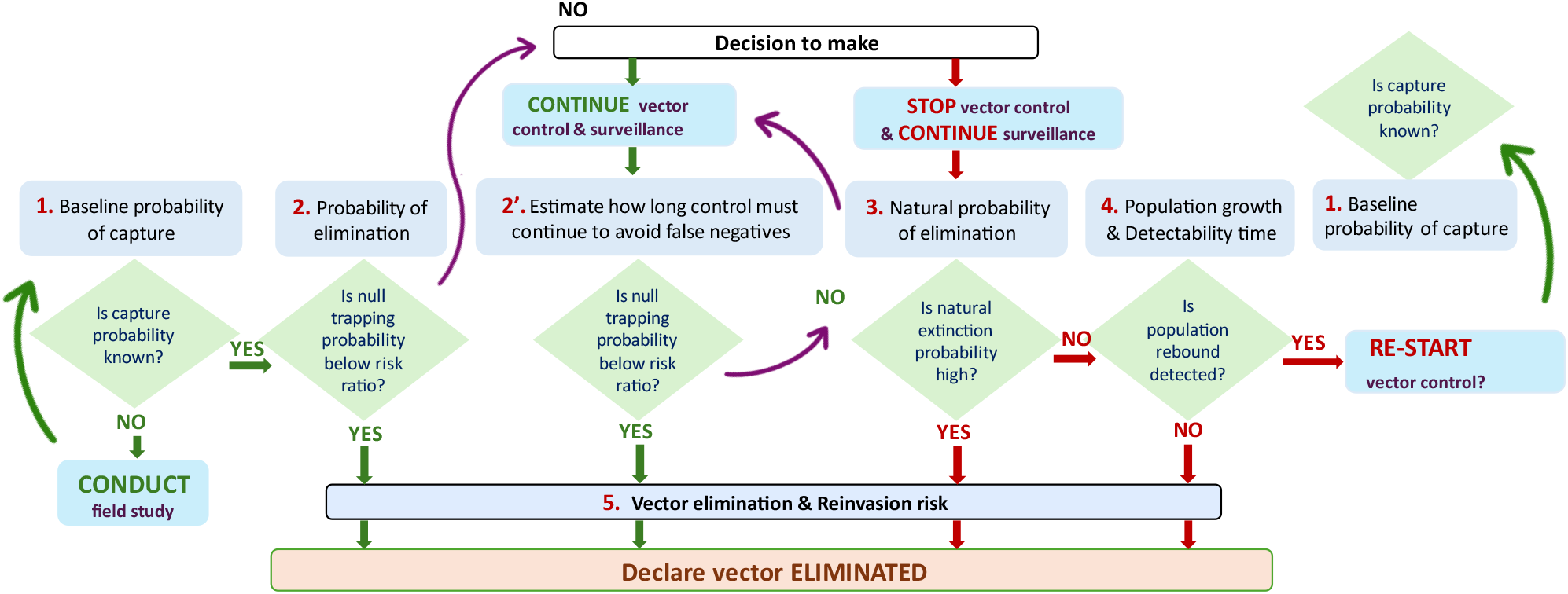
Modelling framework and decision tree for declaring vector elimination: Step 1. Baseline work to assess the probability of vector capture (when we mean known from experiments or estimated from the literature); Step 2. Calculate the conditional probability of observing a series of zero catches given that there is still at least one individual vector present. If that probability is sufficiently low, we conclude the vector has been eliminated (one only enters step 2 once they start having zero catches of vectors in the surveillance system); alternatively step 2’, calculate how long vector control, and surveillance, should continue in order that the risk of obtaining a false negative is acceptably small; Step 3. If the vector population is very low, even if not yet eliminated, calculate the probability of the remnant vector population being eliminated naturally, purely by chance, without further control efforts; Step 4. Use growth models to estimate the expected vector population at various times after the cessation of control efforts, assuming the survival of at least one reproductive female. Failure to detect a rebound in the vector population after protracted periods supports a conclusion that the vector has already been eliminated. If no rebound is detected then the control team must decide whether vector control should resume or whether other disease control actions are necessary (depends of hosts cases surveillance, disease control status and logistical and budgetary constraints); Step 5. Vector elimination can then be declared with a specified risk, and the risk of tsetse reinvasion can also be evaluated.

1. Baseline probability of capture with a surveillance tool: if there is no relevant existing literature conduct mark-recapture, or other, studies to calculate the probability of capturing a vector. Alternatively, calculate the theoretical probability that a given control method could eliminate a vector population that is isolated (*i*.*e*., closed to all in- and out-migration) (32) and infer the probability of capture with the surveillance tool from the vector control tool efficacy.
2. Probabilistic modelling of elimination: apply a probability model to the results of trapping efforts to reject the null hypothesis that vectors are still present in the area.
3. Natural probability of elimination: estimate the probability that a very small residual population will be eliminated by chance.
4. Probability of detecting a rebound: use a model of tsetse population growth to estimate the time required for a very small remnant population to become detectable by trapping.
5. Vector elimination and reinvasion risk: finally, evaluate whether the vector population has been successfully eliminated with a specified level of risk, while also accounting for the potential threat of reinvasion.

Our modelling framework, and the associated decision tree (Figure 8), involves calculating mathematically whether the observed vector surveillance data support a conclusion that elimination has been achieved, what risk would be associated with that conclusion, and the best course of action to progress to elimination.

### Applying the novel vector elimination modelling framework to our case study in Mandoul, Chad

#### Study area

Previously described by Mahamat et al. (13), the Mandoul region covers about 840 km^2^ in parts of five cantons in Southern Chad (Figure 9). It is an historical focus of g-HAT due to *T. brucei gambiense*, transmitted in the area by only one species of tsetse, *G. f. fuscipes*. The mean elevation is ∼400 meters and annual rainfall is between 1000 and 1200 mm: a wet season lasts from June to October and a dry season from November to May. Vegetation, consisting of woody savannah with gallery forest along the rivers, has been degraded in parts through agriculture. The population comprises pastoralist livestock keepers and sedentary mixed crop-livestock farmers – cultivating sorghum, sesame and sweet potatoes.

**Figure 9.**
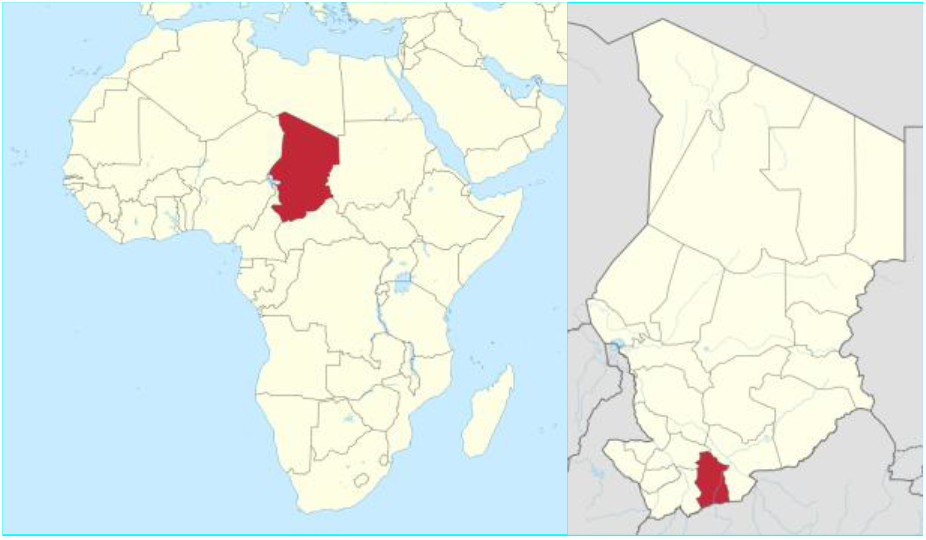
Map of Mandoul area (Chad within the continent, on the left, and Mandoul within Chad on the right).

#### Vector control (Tiny Target deployment)

Recent efforts to eliminate the Mandoul focus of g-HAT are based on the use of Tiny Targets (12) (Figure 10), deployed after a baseline trapping survey had determined the numbers, species and distribution of tsetse, thereby delineating the area to be controlled (Mahamat et al., 2017). The Tiny Targets, provided by Vestergaard (Lausanne, Switzerland), comprised 0.25m × 0.25m blue polyester flanked by 0.25m × 0.25m black polyethylene netting impregnated with deltamethrin at 300mg/m^2^ (12). Targets were deployed along the three main arms of the Mandoul River, where tsetse were detected, and over an area up to 4 km beyond where tsetse were caught. Targets were suspended from tree branches at 10-20 cm above the ground, using string, or erected with wooden sticks obtained locally (Figure 10).

**Figure 10.**
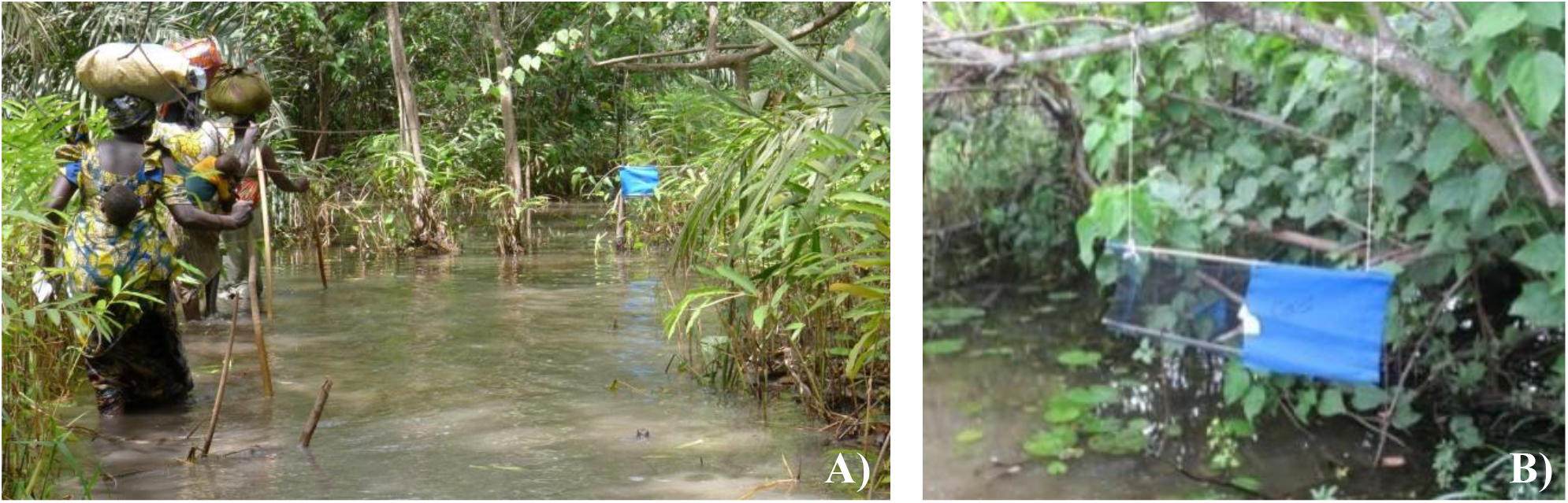
A Tiny Target – as deployed in the Mandoul focus (A: overall view on the river; B: zoom on a target)

A total of 2713 targets were deployed in January-February 2014 and replaced annually until 2022, whereupon 10% of them were removed in 2023, a further 40% removed in 2024, and the last targets removed in April 2025.

#### Vector surveillance

In November 2013, prior to target deployment, the above-cited baseline survey involved the deployment of 108 biconical traps (44) across the whole Mandoul area (Figure 11). The traps were left *in situ* for 48 hours, and the numbers of *G. f. fuscipes* captured in each trap were then recorded. Thereafter, sampling traps were deployed only at 44 sentinel sites and were operated for two days, twice each year, from 2014 to 2024.

**Figure 11.**
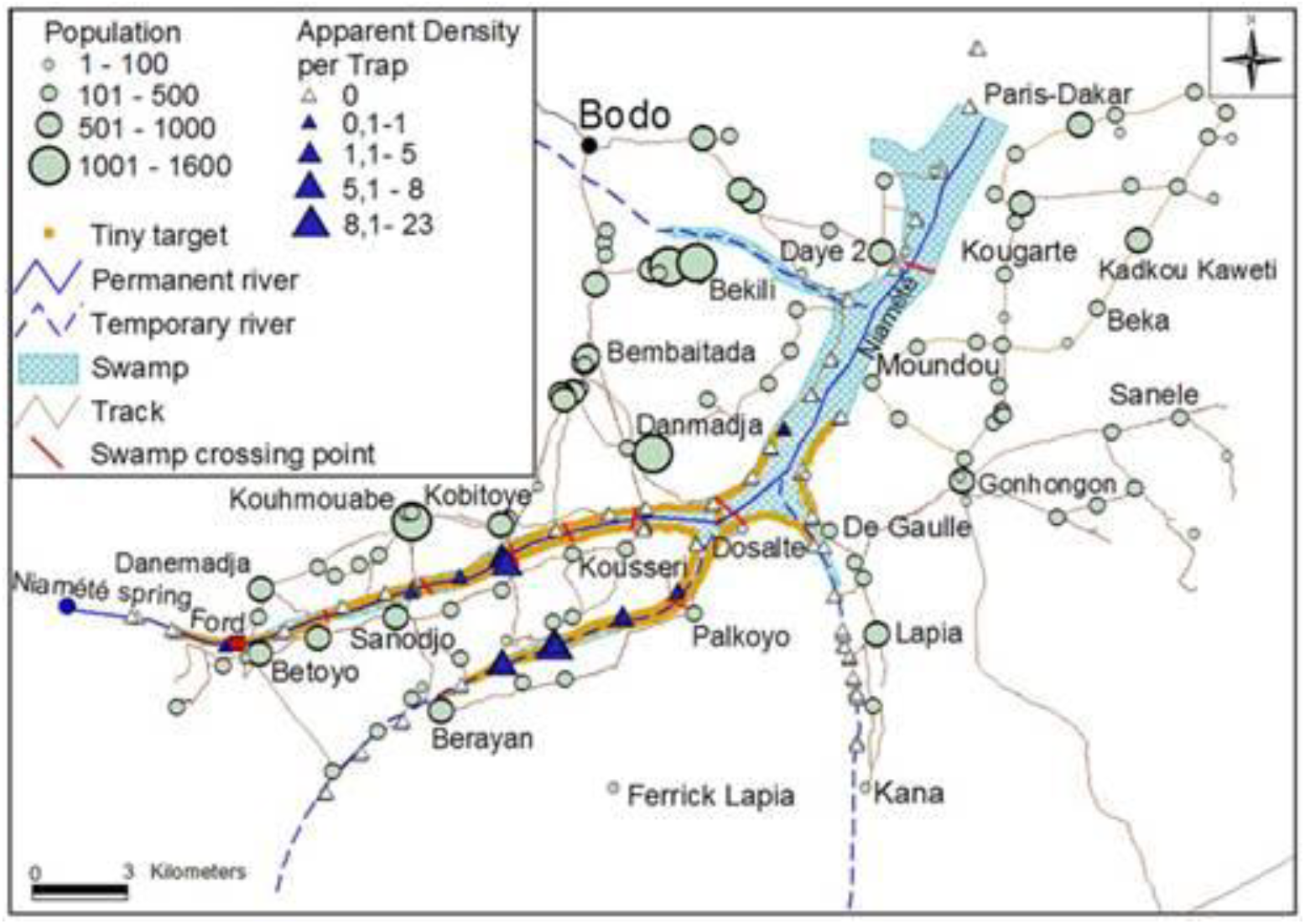
Map showing the deployment positions of the 108 traps used in the preliminary sampling exercise in Mandoul

Tiny targets are used to kill tsetse, whereas biconical traps are only used to catch live tsetse as a monitoring system. Hence, from 2014-2025, tiny targets stayed active all year long (albeit being sometimes replaced by new ones) while traps were only deployed two days at a time, a few times a year, and then removed.

#### The five-step approach in Mandoul

##### Step 1) Baseline probability of capture

Although excellent baseline work, including a comprehensive baseline survey, was carried out in Mandoul, this was a control operation, not a research study – and no trials were carried out to estimate the efficacy of the traps and targets used in the control exercise. Moreover, neither life history nor population dynamics studies were carried out on the population of *G. f. fuscipes* in the Mandoul area. Accordingly, as detailed below, we rely on theory and past examples from other study sites to parametrize some of our models to obtain the probability of capturing a tsetse with the biconical traps used in Mandoul.

We first estimate the theoretical probability that a given vector control method could, in theory, successfully eliminate an isolated population of tsetse (32). We then estimate the probabilities of kill/capture after deployment of targets/traps for a series of days in Mandoul. Finally, using information from Big Chamaunga Island, Kenya, we estimate the probability *p* that a female *G. f. fuscipes*, alive in the Mandoul at the start of a given day, is killed by a target or captured by an individual biconical trap. We assume that we are indeed dealing with an isolated population in Mandoul, since the nearest tsetse population sampled beyond the borders of the Mandoul focus is ∼50km distant (45).

Hargrove (2005) (22) calculated the probability (*s*) of eliminating an isolated tsetse population, as a function of the variables impacting female birth and death rates. The probability is given by the solution of the equation:

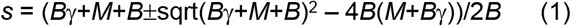

where, for compactness, we write

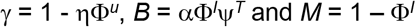

Φ Daily survival probability for adult females.

Ψ Daily survival probability for female pupae.

*u* Period between female adult eclosion and first ovulation (days).

*I* Inter-larval period (days).

*T* Pupal duration (days).

α Probability deposited pupa is female.

*η* Probability adult female is inseminated.

The probability of elimination is the smaller of the two roots of equation (1). This gives the probability that the line emanating from a single female fly is eliminated. If generation zero consists of *N* flies, all subject to the same survival probabilities and reproductive rates, the whole population is eliminated with probability *s*^*N*^.

Evaluation of equation (1) shows that, if the mortality of adult female tsetse in an isolated population can be maintained at a level of at least 3.5 – 4.0% per day, that population will be eliminated with probability 1.0 (Figure 12) – even if there are no reproductive losses, and regardless of density dependent effects (22). The minimum mortality required to be sustained among adult females, in order to achieve elimination, naturally decreases as pupal mortality increases. Thus, if pupal mortality is of the order of at least 1% per day, as estimated for *G. pallidipes* in Kenya (46), a sustained adult mortality of between 2.5 and 3.0% per day will ensure elimination (Figure 12).

**Figure 12.**
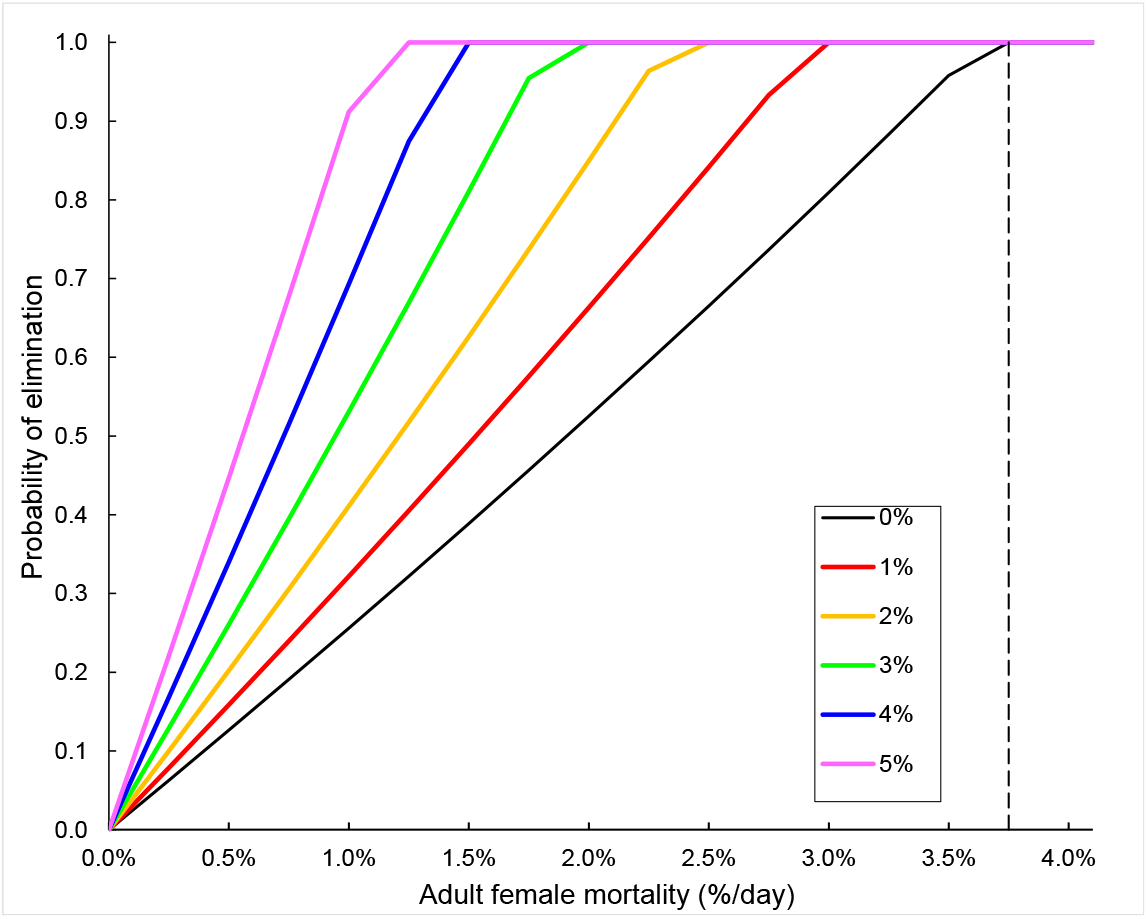
Probability of elimination of an isolated tsetse population as a function of the mortality of female adult females and pupae (legend from 0 to 5%). Calculated for values of u = 7 days, I = 9 days, and T = 27 days. Redrawn from “Hargrove J.W. (2005) Extinction probabilities and times to extinction for populations of tsetse flies Glossina spp (Diptera: Glossinidae) subjected to various control measures. Bulletin of Entomological Research 95, 13-21”.

### Probabilities of kill/capture after deployment of targets/traps for a series of days in Mandoul

For the Chad case study, assume that there is a probability *p* that a female *G. f. fuscipes* alive in the Mandoul at the start of a given day is killed by a target during that day. Then the probability that this fly is *not* killed by a target during that day is *q* = 1 – *p*. In what follows we make frequent use of the assumption that the probability that a fly is *not* killed by any target on a given day is independent of the probability that it was *not* killed by any target on the previous day. By extension, this assumption means that the fly escapes being killed by a target on *n* consecutive days with probability *Q* = *q*^*n*^ – and the probability that the fly has been killed by day *n* is *P* = 1 – *Q* = 1 - *q*^*n*^. An analogous argument applies to the probabilities that a fly is captured, or evades capture, by any given trap.

### Estimate probability p that a female G. f. fuscipes, alive in the Mandoul at the start of a given day, is killed by a target/captured by an individual trap – using information from Big Chamaunga Island, Kenya

The probabilities of trapping *G. f. fuscipes*, or killing them using Tiny Targets, were estimated using data from Tirados et al. (14), who monitored the decline in trap catches of *G. f. fuscipes*, following the deployment of Tiny Targets on Big Chamaunga Island, (−0.426° latitude, 34.233° longitude; surface area 0.2km^2^; circumference 1.5 km), which lies in the Kenyan section of Lake Victoria. From January 2011 - December 2012, they deployed 30 Tiny Targets at 50m intervals in the shoreline habitat of the island, giving a target density of 20 targets/km. They monitored the impact of deploying targets along the island shore via monthly catches of tsetse, from four biconical traps deployed, between 200 and 300m apart, along the lakeshore, and a further single trap placed at the centre of the island.

#### Step 2) probability of elimination based on vector capture

We apply a probability model to the results of our surveillance efforts to reject the null hypothesis that insects are still present following a series of days in Mandoul with no tsetse being captured.

We define:

*A* Area sampled (km^2^), assumed isolated (closed to immigration and emigration).

*N* Total insects surviving the eradication attempt, assumed randomly distributed in *A*.

*σ* Trap efficiency, *i*.*e*., the conditional probability that an insect is caught by a given trap, given that there is only one trap present in the 1-km^2^ square containing the insect and given that the insect is active.

*S* Number of traps present in all of *A*.

*t* Number of days for which each trap is operated.

With these definitions Hargrove (32) showed that the probability of capturing at least one fly is approximately:

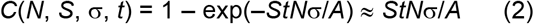

the approximation holding for populations close to elimination, when the exponent is small.

Hence the probability (C’) of null trapping, *i*.*e*., of catching no tsetse at all, is

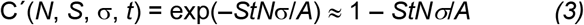

We aim to minimize the risk (α) of falsely concluding – from a sequence of zero catches – that tsetse have been eliminated when, in fact, there are still tsetse present. Accordingly, we typically set α to some value close to zero; say, α = 0.01 or α = 0.001. Then, if we find that C’ ≈ 1 – *StNσ*/*A* < α, we conclude that tsetse have been eliminated, with the attendant risk, α, that our conclusion is false.

#### Step 3) Probability of natural elimination

We estimate the probability that a very small residual population will be eliminated by chance. If only a small number of tsetse survive a control operation, there is a non-zero probability that this remnant population will be eliminated by chance – without the need for further control efforts. The probability that this will occur is calculated using Equation (1) – with the assumption that mortalities among adult and immature females can be much lower than when the population is subjected to control measures. Figure 13 shows that even if only one inseminated female survives, and if the background adult female mortality is 2% per day there is still a 40% chance that the female will give rise to a surviving population. If there are 10 surviving inseminated females, the population will be almost certain to survive – even if adult female mortality is 2.5% per day.

**Figure 13.**
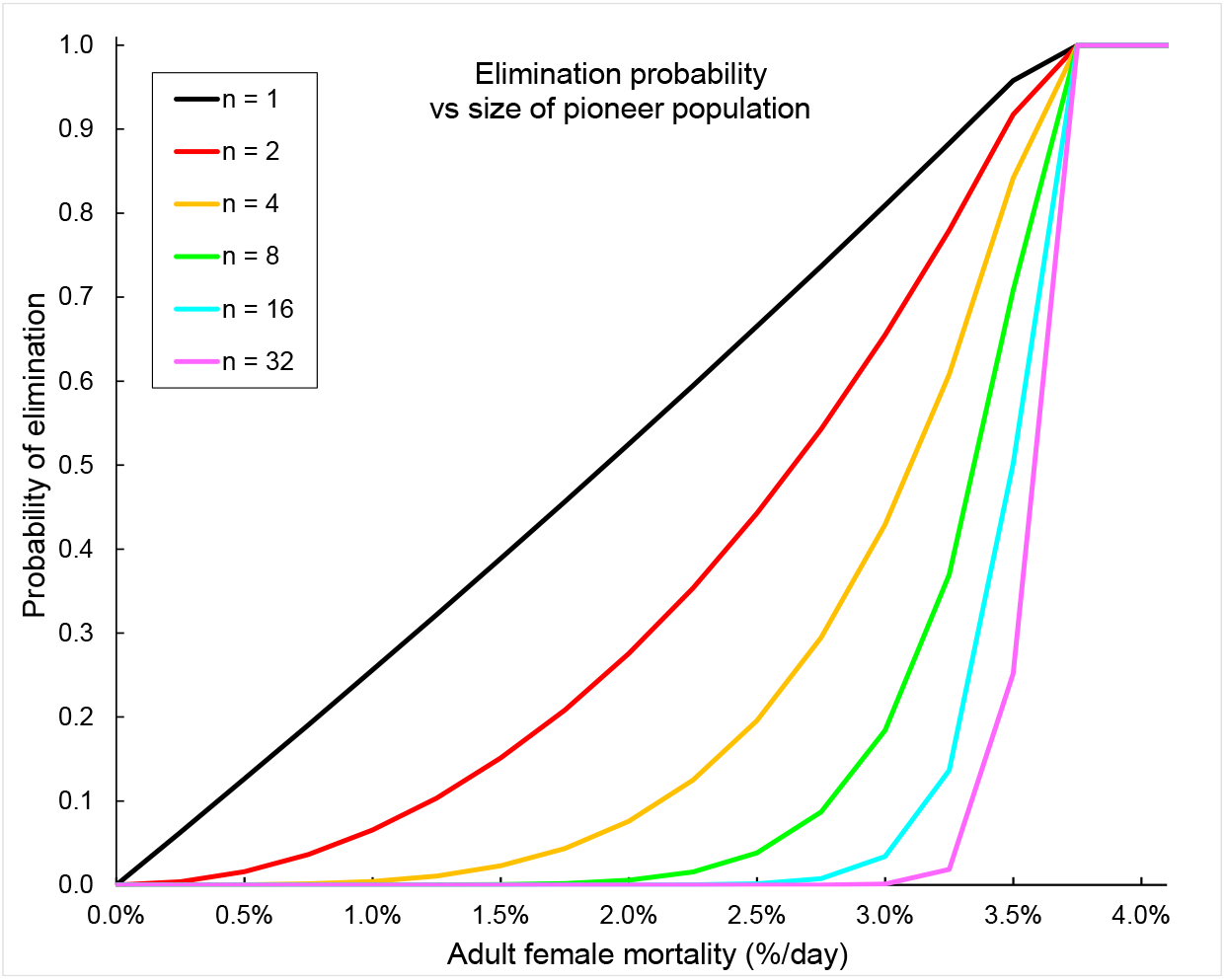
Probability that a small residual tsetse population is eliminated by chance, as a function of adult female mortality and the initial number of females in the residual population. Calculated from Equation (1) with the following input parameters: Time to first ovulation, u=7 days; Inter-larval period, I=9 days. Pupal duration, T=30 days; Probability deposited pupa is female, α=0.5; Probability female is inseminated, η=1.0. Redrawn from “Hargrove J.W. (2005) Extinction probabilities and times to extinction for populations of tsetse flies Glossina spp (Diptera: Glossinidae) subjected to various control measures. Bulletin of Entomological Research 95, 13-21”.

#### Step 4) Detection of potential rebound

If there is insufficient evidence – even after a series of zero catches of tsetse – to conclude that tsetse have been eliminated, or that the remaining small tsetse population may be eliminated naturally, then there are two options.

The first, which we call “ step 2’ “ is to keep the vector control/Tiny Targets in place and to keep sampling with traps until the sequence of zero catches is so long that we can have a high degree of confidence that elimination has been achieved. Hence, using Equation 2, one calculates the sampling duration needed before reaching a high confidence (below the risk ratio) that the vector population has been eliminated. If feasible one could calculate a combination of increased sampling efforts (more traps) and prolonged surveillance efforts. This has the advantage that the presence of the Tiny Targets will ensure a low risk of a recurrence of cases of human trypanosomiasis. It has the disadvantage, however, of incurring, for an unknown/long period, the continued costs of employing the control team, and buying and deploying new Tiny Targets. Moreover, these costs are wasted if the tsetse population has, in fact, already been eliminated.

The alternative is to remove all Tiny Targets from the control area, here Mandoul, and to continue with the vector sampling effort – but to stop all control measures and aim at detecting a potential rebound in vector population, which is our actual step 4. There are two possible outcomes of step 4: (i) The tsetse population has indeed already been eliminated – in which case the ongoing sampling will fail to catch a fly, regardless of how long the sampling continues: there is then no longer any need to carry out any manner of vector or disease control. (ii) The more interesting, and problematic, possibility is that the surviving tsetse are not eliminated by chance after the Tiny Targets have been removed. If this is the case, we expect the tsetse population to grow steadily, particularly given that there is no longer any risk of the flies being killed by Tiny Targets. Then one has to determine whether vector control should be restarted.

Failure to detect a rebound in the vector population after the cessation of control efforts will support the conclusion that the vector has already been eliminated. A growth model is used to estimate the expected vector population at various times after the cessation of control efforts, assuming the survival of at least one reproductive female. If no tsetse can be captured, despite predictions of a large population from the growth model, then the vector population can be considered as eliminated.

The growth of an isolated population of tsetse is determined by the balance between the rates of larval production, and development – and by the rates of immature and adult mortality, whether natural or imposed by human intervention. Hargrove (33) estimated growth rates of tsetse populations, as functions of birth and death rates, by calculating dominant eigenvalues of appropriate Leslie matrices. Those results, summarized in Figure 5, are valid for any tsetse population; it is only necessary to stipulate the appropriate levels of the birth and death rates. In setting development rates for tsetse in the Mandoul area, we assume a mean daily temperature of 28.4°C, based on its hot dry equatorial location. In the absence of field estimates of the effects of temperature on various development rates in *G. f. fuscipes*, we use relationships measured for *G. m. morsitans* and *G. pallidipes*, which deposit their first larva at the age of about 14 days, and subsequent larvae at 8-day intervals (32, 47, 48). Pupal duration for female *G. m. morsitans*, at a constant temperature of 28.4°C in the laboratory, is 21 days (49, 50). In the field, however, temperatures in typical larviposition sites are about 2°C cooler on average than ambient (51, 52). Accordingly, we assume a temperature of 26.4°C during pupal development, giving an expected pupal duration of 24 days. We estimate possible growth rates for tsetse populations in the Mandoul area, before and after the use of Tiny Targets, for a wide range of adult mortalities (Figure 13).

Having predicted the growth of the tsetse population, one can then use Equations (2) and (3) of step 2 to determine when we will have a 90%, 99% or 99.9% confidence that our zero catches, and failure to detect a rebound, actually means elimination.

#### Step 5) Vector elimination and reinvasion risk

Finally, we evaluate whether the vector population has been successfully eliminated with a specified level of risk, while also accounting for the potential threat of reinvasion. The above methodologies, for estimating the probability of elimination of a tsetse population, apply to situations where the tsetse population is isolated. If, however, we conclude that tsetse have been eliminated from the Mandoul, we still need to estimate the probability that the area could be repopulated by tsetse invading from distant populations. The nearest tsetse population, beyond the borders of the Mandoul focus is ∼50km distant (45).

Dispersal in tsetse is generally modelled as a random walk (53–55) or, equivalently, as a diffusion process (27). This last paper was published only as a hard copy and is not generally available. Accordingly, it is included here as Supplementary file S1 and is used to estimate the probability that a tsetse fly, present at time 0 at a random point in a given area, will be found in some distant neighborhood at time *t* later, given that it is still alive. In making these calculations for *G. f. fuscipes*, we use – as a first approximation – Rogers’ (54) estimate that *G. f. fuscipes* moves an average of 137 m (150 yds) per day and assume that this rate of movement does not vary significantly with age.

The closest population of tsetse, *G. f. fuscipes*, beyond the borders of the Mandoul focus is in the neighbourhood of Timbéri, some 50 km distant. We estimate the probability that a tsetse fly could survive long enough to move between the two populations. We use the results of Hargrove & Lange (27) in modelling tsetse dispersal as diffusion in the plane, starting at the origin when time *t* = 0, with coefficient of diffusion σ^2^. The fly’s position (*x, y, t*) in time and space is then defined by a normally distributed random variable with density function

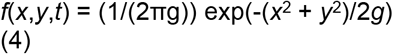

where

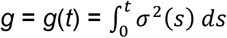

As a first approximation we assume σ^2^ is independent of the fly’s age and position in the plane, so that *g* = *kt*, where *k* is a constant. Consider a fly starting its dispersal at a point chosen uniformly from the interval [*a, b*]. Then at some time *t* it will be in the interval [*c, d*] with probability:

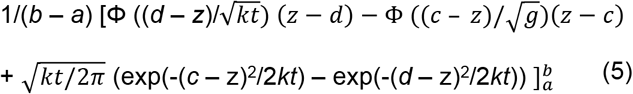

Hargrove & Lange (1989) note that if the intervals [*a, b*] and [c, d] are widely separated, relative to the rate of diffusion, such that (c-a)/(*kt*)^0.5^ >> 0, then Equation (5) can be much simplified because:

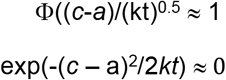

## Supporting information

Supplementary file S1 (Hargrove & Lange, 1989) paper

## Acknowledgments

Dedicated to the memories of Hugh Barclay (1941 – 2022), who pioneered efforts to support the declaration of elimination of tsetse populations, of Jean-Baptiste Rayaisse (1967-2020) who was a fantastic tsetse entomologist, a hard worker, deeply committed to trypanosomiasis control, and with a great sense of humour, and to Ali Bachar Alkatib, killed by bees during the first target deployment in Mandoul. We acknowledge inputs from Profs Glyn Vale and Steve Torr on other case studies and to guide parametrization.

## Author Contributions

**JH** conceptualization, data curation, formal analysis, investigation, methodology, software, validation, visualization, Writing – Original Draft Preparation; **MHM** investigation, validation, resources, Writing – Original Draft Preparation, **MA** investigation, validation, resources, Writing – Original Draft Preparation; **WY** investigation, validation, resources, Writing – Original Draft Preparation, **DS** investigation, validation, Writing – Review & Editing; **JD** investigation, resources, Writing – Review & Editing; **ES** investigation, resources, Resources, Writing – Review & Editing; **IK** Methodology, Resources, visualization, Writing – Review & Editing; **AM** Resources, Writing – Review & Editing; **PB** formal analysis, visualization, Writing – Original Draft Preparation; **PS** conceptualization, Funding Acquisition, Project Administration, methodology, validation, Writing – Original Draft Preparation; **AMGB** conceptualization, data curation, Funding Acquisition, Project Administration, formal analysis, investigation, methodology, software, validation, visualization, Writing – Original Draft Preparation

## Data, Materials, and Software Availability

All necessary data are available within the supplementary material or upon request and all equations are available within the main text.

## Supplementary material

- Supplementary file S1 (Hargrove & Lange, 1989) paper
- Supplementary file S2 processed data tsetse capture baseline Mandoul in 2013 and capture surveys 2014-2024 (available upon request)
- Supplementary file S3 processed data on catches of female *G. f. fuscipes*, on Big Chamaunga Island, Lake Victoria, Kenya (available upon request)

## References

1. World Health Organization, Vector-borne diseases. WHO (2024).

2. J. R. Franco, et al., The elimination of human African trypanosomiasis: Achievements in relation to WHO road map targets for 2020. PLoS Negl. Trop. Dis. 16, e0010047–e0010047 (2022).

3. A. Boisson-Walsh, Chad eliminates gambiense sleeping sickness. Lancet Infect. Dis. 24, e551–e551 (2024).

4. R. Nuwer, Why the last cases of sleeping sickness will be the hardest to eliminate. Nature (2025). 10.1038/D41586-025-00013-6.

5. D. Kaba, et al., Towards the sustainable elimination of gambiense human African trypanosomiasis in Côte d’Ivoire using an integrated approach. PLoS Negl. Trop. Dis. 17, e0011514–e0011514 (2023).

6. J. R. Franco, et al., The elimination of human African trypanosomiasis: Monitoring progress towards the 2021–2030 WHO road map targets. PLoS Negl. Trop. Dis. 18 (2024).

7. P. Barreaux, et al., Priorities for Broadening the Malaria Vector Control Tool Kit. Trends Parasitol. 33, 763–774 (2017).

8. P. A. Hancock, et al., Mapping trends in insecticide resistance phenotypes in African malaria vectors. PLoS Biol. 18 (2020).

9. J. Hemingway, et al., Tools and Strategies for Malaria Control and Elimination: What Do We Need to Achieve a Grand Convergence in Malaria? PLoS Biol. 14 (2016).

10. A. F. Read, P. A. Lynch, M. B. Thomas, How to make evolution-proof insecticides for malaria control. PLoS Biol. 7, e1000058–e1000058 (2009).

11. A. M. G. Barreaux, et al., The role of human and mosquito behaviour in the efficacy of a house-based intervention. Philos. Trans. R. Soc. B Biol. Sci. 376, 20190815–20190815 (2021).

12. J. Esterhuizen, et al., Improving the cost-effectiveness of visual devices for the control of riverine tsetse flies, the major vectors of human African trypanosomiasis. PLoS Negl. Trop. Dis. 5 (2011).

13. M. H. Mahamat, et al., Adding tsetse control to medical activities contributes to decreasing transmission of sleeping sickness in the Mandoul focus (Chad). PLoS Negl. Trop. Dis. 11, e0005792–e0005792 (2017).

14. I. Tirados, et al., Tsetse control and Gambian sleeping sickness; implications for control strategy. PLoS Negl. Trop. Dis. (2015). 10.1371/journal.pntd.0003822.

15. K. S. Rock, et al., Update of transmission modelling and projections of gambiense human African trypanosomiasis in the Mandoul focus, Chad. Infect. Dis. Poverty 2022 111 11, 1–13 (2022).

16. S. English, et al., “Incorporating Vector Ecology and Life History into Disease Transmission Models: Insights from Tsetse (Glossina spp.)” in Population Biology of Vector-Borne Diseases, Oxford University Press, 2020), pp. 175–188.

17. A. M. G. Barreaux, P. Barreaux, K. Thievent, J. C. Koella, Larval environment influences vector competence of the malaria mosquito Anopheles gambiae. MalariaWorld J. 7 (2016).

18. A. M. G. Barreaux, C. M. Stone, P. Barreaux, J. C. Koella, The relationship between size and longevity of the malaria vector Anopheles gambiae (s.s.) depends on the larval environment. Parasit. Vectors 11, 485–485 (2018).

19. A. J. Mackay, J. Yan, C. H. Kim, A. M. G. Barreaux, C. M. Stone, Larval diet and temperature alter mosquito immunity and development: using body size and developmental traits to track carry-over effects on longevity. Parasit. Vectors 16, 1–13 (2023).

20. J. S. Lord, J. W. Hargrove, S. J. Torr, G. A. Vale, Climate change and African trypanosomiasis vector populations in Zimbabwe’s Zambezi Valley: A mathematical modelling study. PLOS Med. 15, e1002675–e1002675 (2018).

21. J. S. Lord, et al., Effects of maternal age and stress on offspring quality in a viviparous fly. Ecol. Lett. ele.13839-ele.13839 (2021). 10.1111/ELE.13839.

22. J. W. Hargrove, Extinction probabilities and times to extinction for populations of tsetse flies Glossina spp. (Diptera: Glossinidae) subjected to various control measures. Bull. Entomol. Res. 95, 13–21 (2005).

23. H. J. Barclay, J. W. Hargrove, Probability models to facilitate a declaration of pest-free status, with special reference to tsetse (Diptera: Glossinidae). Bull. Entomol. Res. 95, 1–11 (2005).

24. E. B. Are, J. W. Hargrove, J. Dushoff, Does Counting Different Life Stages Impact Estimates for Extinction Probabilities for Tsetse (Glossina spp)? Bull. Math. Biol. 83, 1–13 (2021).

25. G. A. F. Seber, The estimation of animal abundance : and related parameters, Second edition. (Griffin, 1982).

26. B. K. Williams, J. D. Nichols, M. J. Conroy, Analysis and Management of Animal Populations: Modeling, Estimation and Decision Making. (2002).

27. J. W. Hargrove, K. Lange, Tsetse dispersal viewed as a diffusion process. Trans. Zimb. Sci. Assoc. 64, 1–8 (1989).

28. L. R. Haines, et al., Big Baby, Little Mother: Tsetse Flies Are Exceptions to the Juvenile Small Size Principle. BioEssays 2000049–2000049 (2020). 10.1002/bies.202000049.

29. J. W. Hargrove, M. O. Muzari, S. English, How maternal investment varies with environmental factors and the age and physiological state of wild tsetse Glossina pallidipes and Glossina morsitans morsitans. R. Soc. Open Sci. 5, 171739–171739 (2018).

30. K. Heller, Tsetse Flies Rely on Symbiotic Wigglesworthia for Immune System Development. PLoS Biol. 9, e1001070–e1001070 (2011).

31. B. L. Weiss, J. Wang, S. Aksoy, Tsetse Immune System Maturation Requires the Presence of Obligate Symbionts in Larvae. PLOS Biol. 9, e1000619–e1000619 (2011).

32. J. W. Hargrove, Tsetse eradication: sufficiency, necessity and desirability Telephone. DFID Anim. Health Programme 133-+ix (2003).

33. J. W. Hagrove, Tsetse: the limits to population growth. Med. Vet. Entomol. 2, 203–217 (1988).

34. D. A. Turner, R. Brightwell, An evaluation of a sequential aerial spraying operation against Glossina pallidipes Austen (Diptera: Glossinidae) in the Lambwe Valley of Kenya: aspects of post-spray recovery and evidence of natural population regulation. Bull. Entomol. Res. 76, 331–349 (1986).

35. G. A. Vale, J. W. Hargrove, G. F. Cockbill, R. J. Phelps, Field trials of baits to control populations of Glossina morsitans morsitans Westwood and G. pallidipes Austen (Diptera: Glossinidae). Bull. Entomol. Res. 76, 179–179 (1986).

36. R. D. Dransfield, The Range of Attraction of the Biconical Trap for Glossina pallidipes and Glossina brevipalpis. Int. J. Trop. Insect Sci. 1984 55 5, 363–368 (1984).

37. P. M. Kgori, S. Modo, S. J. Torr, The use of aerial spraying to eliminate tsetse from the Okavango Delta of Botswana. Acta Trop. 99, 184–199 (2006).

38. J. W. W. S., The eradication of sleeping sickness from Principe. Nature 98, 311–312 (1917).

39. R. Du Toit, Trypanosomiasis in Zululand and the control of tsetse flies by chemical means. (1954).

40. M. J. B. Vreysen, et al., Glossina austeni (Diptera: Glossinidae) Eradicated on the Island of Unguja, Zanzibar, Using the Sterile Insect Technique. J. Econ. Entomol. 93, 123–135 (2000).

41. J. Bouyer, et al., Mapping landscape friction to locate isolated tsetse populations that are candidates for elimination. Proc. Natl. Acad. Sci. U. S. A. 112, 14575–14580 (2015).

42. A. H. Dicko, et al., Using species distribution models to optimize vector control in the framework of the tsetse eradication campaign in Senegal. Proc. Natl. Acad. Sci. 111, 10149–10154 (2014).

43. O. Diall, et al., Developing a Progressive Control Pathway for African Animal Trypanosomosis. Trends Parasitol. 33, 499–509 (2017).

44. A. Challier, C. Laveissière, Un nouveau piège pour la capture des glossines (Glossina: Diptera, Muscidae) déscription et essais sur la terrain. Cah. ORSTOM 11, 251–262 (1973).

45. S. Ravel, et al., Population genetics of Glossina fuscipes fuscipes from southern Chad. Peer Community J. 3, e31–e31 (2023).

46. D. J. Rogers, S. E. Randolph, Estimation of rates of predation on tsetse. Med. Vet. Entomol. 4, 195–204 (1990).

47. J. W. Hargrove, Reproductive rates of tsetse flies in the field in Zimbabwe. Physiol. Entomol. 19, 307–318 (1994).

48. J. W. Hargrove, Towards a general rule for estimating the stage of pregnancy in field-caught tsetse flies. Physiol. Entomol. 20, 213–223 (1995).

49. R. J. Phelps, P. M. Burrows, Puparial duration in glossina morsitans orientalis under conditions of constant temperature. Entomol. Exp. Appl. 12, 33–43 (1969).

50. J. W. Hargrove, G. A. Vale, Models for the rates of pupal development, fat consumption and mortality in tsetse (Glossina spp). Bull. Entomol. Res. 1–13 (2019). 10.1017/S0007485319000233.

51. R. J. Phelps, P. M. Burrows, Lethal temperatures for puparia of glossina morsitans orientalis. Entomol. Exp. Appl. 12, 23–32 (1969).

52. P. J. Jackson, R. J. Phelps, Temperature regimes in pupation sites of Glossina morsitans orientalis Vanderplank (Diptera). Rhod. Zamb. Malawi J. Agric. Res. 5, 249–260 (1967).

53. E. Bursell, “Dispersal and concentration of Glossina” in The African Trypanosomiases, H. W. Mulligan, Ed. (George Allen and Unwin, 1970), pp. 382–394.

54. D. Rogers, Study of a Natural Population of Glossina fuscipes fuscipes Newstead and a Model of Fly Movement. J. Anim. Ecol. 46, 309–309 (1977).

55. J. W. Hargrove, Tsetse Dispersal Reconsidered. J. Anim. Ecol. 50, 351–351 (1981).

