## Supplementary file S1 (Hargrove & Lange, 1989) paper for "Modelling framework to demonstrate elimination of a vector population: tsetse elimination in Chad"

### Tsetse dispersal viewed as a diffusion process

**J. W. Hargrove and K. Lange**

*Department of Biomathematics*

*University of California. Los Angeles, California 90024, U.S.A*

#### SUMMARY

Tsetse dispersal is viewed as a diffusion process, with the position of a fly, relative to its origin, a normally distributed random variable. Defining $R$ as the mean distance of a diffusing particle from the origin and $R_d = R$ for $0 \leq R \leq d$, and 0 otherwise, the moments of $R_d$ are calculated. These are used, in turn, to compute the conditional moments $(E(R^k|R \leq d)$; together these define the diffusion of the particle out of a circular domain. Similar results are produced for diffusion out of a rectangle and into adjoining rectangles. The results are used to model the dispersal of tsetse flies as measured in mark-recapture experiments; they provide a reasonable description of Jackson's (1946) classical 'square spiral' data, but his 'large square' data do not conform well to the simplest diffusion model. An age dependent coefficient of diffusion, or a prolonged 'escape reaction', are possible reasons which would need to be invoked unless more complex models of movement are involved. The classical data do not allow us to separate between these possibilities.

#### INTRODUCTION

It has been suggested that dispersal in tsetse (*Glossina* spp) can be viewed as a series of discrete daily steps each taken in a random direction (Bursell, 1970; Rogers, 1977). Hargrove (1981) suggested an elaboration of this model whereby the step length may vary and, in particular, probably change with the age and physiological state of the fly.

In the present work tsetse dispersal is viewed as a diffusion process, which is a very similar process to those considered previously; the 'rate' of dispersal is simply defined as a diffusion coefficient rather than as a discrete step length. A number of results, defining the spatio-temporal distribution of the diffusing particle, then emerge which are not readily available via the discrete method (Skellam, 1951). Diffusion models have been used to estimate the rates of movement of several insects (see, for example, Dobzhansky & Wright, 1943; Crumpacker and Williams, 1973; Kareiva, 1983). Although the model is only a first approximation to the true situation, and the underlying biological assumption of lack of interaction of individuals is biologically unsound (Taylor, 1980), it does have the advantage of affording easy statistical estimation of a measure of the rate of dispersal, and other allied parameters, and will be sufficiently accurate in many cases for the practical biologist. It has the advantage over the discrete-step random walk model, of allowing one to model the results of relevant experiments using regression techniques, rather than simulation (*cf* Bursell, 1970; Hargrove, 1981); and thus to choose between competing models using objective statistical criteria.

#### RESULTS AND DISCUSSION

*Theoretical development*

The model considered here is for a diffusion coefficient which may vary with time (normally the age of the fly in the case of tsetse), but is independent of position. If a particle moves by diffusion in the plane, starting at the origin when time $t = 0$, with coefficient of diffusion $\sigma^2(t)$ then the position of the particle in time and space is defined by a normally distributed random variable with density function

$$f(x,y,t) = (1/(2\pi g)) \exp(-(x^2 + y^2)/2g) \quad (1)$$

where $$g = g(t) = \int_0^t \sigma^2(s)\, ds$$

(see Kannan (1979) for an example of the derivation of this equation). In words, $f(x, y, t)$ is the probability per unit area that the diffusing particle will be found around the point $(x, y)$ at time $t$.

To find the probability of the particle being in some particular region of the plane at a given time one must integrate $f(x,y,t)$ over the region. In general this is not easy, but for shapes such as circles and rectangles the problem is simplified and, for our purposes, these results. are quite adequate.

*Diffusion in a circular domain*

We consider first diffusion starting at the centre of a circular domain. By symmetry the mean displacement of a diffusing particle from its starting point is zero for all time, but it is of interest to investigate the random distance

$$R = \sqrt{x^2 + y^2}$$

of the particle from the origin as a function of time. Since any experiment is necessarily confined to a finite domain we look at the moments of the truncated distance $R_d$ where $R_d = R$ for $0 \leq R \leq$, d and 0 otherwise, defined for all $d > 0$. By changing Equation 1 to polar coordinates, the moments of R can be expressed as:

$$E(R_d^k) = (1/2\pi g) \int_0^d \int_0^{2\pi} r^k \exp(-r^2/2g) r \; d\theta \; dr$$

$$= \int_0^d r^k \exp(-r^2/2g) r/g \; dr \quad (2)$$

for $k = 0, 1, 2 \ldots.$

When $k = 0$ $$E(R_d^0) = 1 - \exp(-d^2/2g) \quad (3)$$

And this is simply the probability ($P(R \leq d)$ that the diffusing particle is within distance $d$ of the origin at time $t$. For $k > 0$ integrate by parts in (2) to achieve the reduction

$$E(R_d^k) = -d^k \exp(-d^2/2g) + \int_0^d k r^{k-1} \exp(-r^2/2g) \; dr \quad (4)$$

When $k = 1$, the integral in (4) becomes

$$\int_0^d k r^{k-1} \exp(-r^2/2g) \; dr = \sqrt{\pi g/2} \; (1 - 2\Phi(-d/\sqrt{g}))$$

where $\Phi$ is the standard normal distribution function

$$\Phi(x) = (1/2\pi) \int_{-\infty}^{x} \exp(-u^2/2)\, du$$

Thus $$E(R_d^1) = \sqrt{\pi g/2}\ (1\text{-}2\Phi(\text{-}d/\sqrt{g}) - d \exp(-d^2/2g) \qquad (5)$$

For $k > 1$, (4) can be written as a recurrence relation:

$$E(R_d^k) = -\, d^k \exp(-d^2/2g) + kg\, E(R_d^{k-2})$$

and evaluated using $E(R_d^0)$ and $E(R_d^1)$ from (3) and (5). For instance,

$$E(R_d^2) = -\, d^2 \exp(-d^2/2g) + 2g\ (1 - \exp(-d^2/2g) \qquad (6)$$

In the limit as $d \Rightarrow \infty$, the non-truncated moments become available. Hence the mean and variance of of the distance ($R$) moved from the origin are given by

$$E(R) = \sqrt{\pi g/2} \qquad (7)$$

$$\mathrm{Var}\,(R) = E(R^2) - (E(R))^2 \qquad (8)$$

$$= g(2 - \pi/2)$$

The truncated moments can also be used to compute the conditional moments $E(R^k\,|R \leq d)$ via

$$E(R^k \mid R \leq d) = E(R_d^k)/E(R_d^0)$$

For example, using (3), (5) and (6) the conditional mean is

$$(E(R \mid R \leq d)) = [\sqrt{\pi g/2}\ (1\text{-}2\Phi(\text{-}d/\sqrt{g})) - d \exp(-d^2/2g)]/[1 - \exp(-d^2/2g)] \qquad (9)$$

and the conditional variance is

$$\mathrm{Var}(R \mid R \leq d) = \frac{2g\,(1-\exp(-d^2/2g))-d^2\exp(-d^2/2g)}{1 - \exp(-d^2/2g)} - (E(R \mid R \leq d))^2 \qquad (10)$$

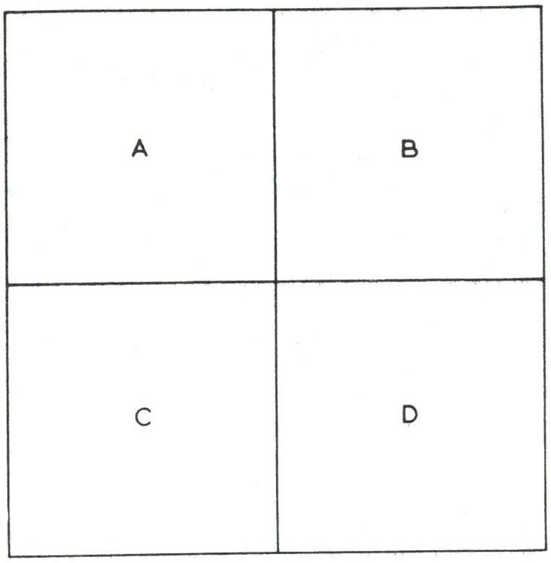


**Figure 1.** Jackson's large square. Tsetse were caught randomly in each of the four sub-squares and marked, with dots of artists' oil paint, to indicate the time and position of their release. Each sub-square had side 2 miles.

The conditional moments are important for real experiments where animals are only sampled at small distances from the origin, relative to the capacity of movement of the animal being studied. Where, as in the experiments of Jackson (1941, 1946) with tsetse, the area is relatively large, (7) and (8) will be adequate approximations to (9) and (10).

*Diffusion in a rectangle*

Jackson (1941) released tsetse in a (4 x 4) mile square of their habitat and observed the relative probability of recapturing flies in the quadrant of their release and in adjacent quadrants (Fig. 1). We consider here the general problem of the probability of a particle, released at time zero at some random point in rectangle A, being present in rectangle B at time $t$. For simplicity we consider the case where A and B have sides parallel to the axes. Since the $X$ position is independent of the $Y$ position the problem really reduces to two one-dimensional problems. The corresponding probabilities are then multiplied to give the final answer.

Thus, consider the one-dimensional $X$ motion of the particle. Suppose the starting point is chosen uniformly from the interval $[a, b]$. Then at some time $t$ it will be in the interval $[c, d]$ with probability

$$1/(b-a) \int_a^b (1/\sqrt{2\pi g}) \int_c^d \exp(-(x-z)^2/2g)\,dx\,dz$$

$$= 1/(b-a) \int_a^b (\Phi((d-z)/\sqrt{g}) - \Phi((c-z)/\sqrt{g}))\,dz \qquad (11)$$

Integrating by parts, changing variables, and applying the Fundamental Theorem of Calculus yields

$$\int_a^b \Phi((d-z)/\sqrt{g})\mathrm{d}z = \Phi((d-z)/\sqrt{g})\,(z-d)|_a^b - \sqrt{g/2\pi}\,\exp(-(d-z)^2/2g)|_a^b$$

The second part of the integral in (11) can be evaluated in like manner, and the relevant probability of being in $[c, d]$ at time t is:

$$1/(b-a)\,[\Phi((d-z)/\sqrt{g})\,(z-d) - \Phi((c-z)/\sqrt{g})(z-c)$$

$$+ \sqrt{g/2\pi}\,(\exp(-(c-z)^2/2g) - \exp(-(d-z)^2/2g))\,]_a^b \qquad (12)$$

In some circumstances this cumbersome expression simplifies. For instance, when $a = c$,

$$\Phi((c-a)/\sqrt{g})(a-c) = 0$$

$$\exp(-(c-a)^2/2g) = 1$$

when $(c-a)/\sqrt{g} \ll 0$

$$\Phi((c-a)/\sqrt{g}) \approx 0$$

$$\exp(-(c-a)^2/2g) \approx 0$$

and when $(c-a)/\sqrt{g} >> 0$

$$\Phi((c-a)/\sqrt{g}) \approx 1$$

$$\exp(-(c-a)^2/2g) \approx 0$$

Application of (12) produces results which are of interest in the analysis of Jackson's data. When the release point is chosen at random in a square $L$ of side $2l = b - a$ (refer to (12) and to Fig. 1) the probability that the particle is still in the large square $L$ after time $t$ is

$$P_L(t) = (\Phi(2l/\sqrt{g}) - \Phi(-2l/\sqrt{g}) - \sqrt{g/2\pi l^2}\,(1 - \exp(-2l^2/g)))^2 \qquad (13)$$

When $l$ is large relative to $\sqrt{g}$, we have the approximation

$$P_L(t) \approx (1 - \sqrt{g/2\pi l^2})^2 \approx 1 - \sqrt{2g/\pi l^2} \qquad (14)$$

Again referring to Fig. 1, if we restrict the random starting points to the small square A (of side $l$), the probabilities that the particle will be in A, B (or equivalently C) or D can similarly be obtained from (12). Approximations for large $l\sqrt{g}$, are given by:

$$P_A(t) \approx 1 - 2\sqrt{2g/\pi l^2} \qquad (15)$$

$$P_B(t) \approx \sqrt{(g/8\pi l^2)}\,(1 - 2\sqrt{2g/\pi l^2}) \qquad (16)$$

$$P_D(t) \approx g/8\pi l^2 \qquad (17)$$

Finally, we consider the simplification where the particle begins diffusing from some fixed point inside a rectangle $r$. (This will be of value in many mark release experiments where animals are released from a single point). The probability ($P_r(t)$ that the particle. will be within $r$ at time $t$ later is given by

$$P_r(t) = (\Phi(l_2/\sqrt{g}) - (\Phi(l_1/\sqrt{g}))(\Phi(l_4/\sqrt{g}) - (\Phi(l_3/\sqrt{g})) \qquad (18)$$

where $l_1$, $l_2$, $l_3$, $l_4$, are the distances of the sides of the rectangle from the starting point of the process. For a square s of side $2l$, (18) becomes

$$P_S(t) = (\Phi(l/\sqrt{g}) - \Phi(-l/\sqrt{g}))^2 \qquad (19)$$

*Regression analyses of data available for tsetse dispersal studies*

Previous studies on tsetse dispersal (Jackson, 1941, 1946; Bursell, 1970; Rogers, 1977; Hargrove, 1981) have used, almost exclusively, the data produced in the 1930s and 1940s by Jackson. There are in fact no other data of comparable scope in the literature. The experiments have been discussed in detail elsewhere (Hargrove, 1981) and will only be described briefly here.

Jackson performed two basic types of experiments on the dispersal of the male tsetse fly *G. morsitans morsitans* Westwood. In the first (referred to here as the 'square spiral' experiment) Jackson (1946) released young flies at the 'centre' of a square spiral path (Fig. 2). Men using hand nets walked the entire spiral each day and caught all available tsetse and, since the design provided approximately equal lengths of path per unit area at all distances from the release point, the distribution of the recaptures can be used to infer rates of dispersal for this species of tsetse.

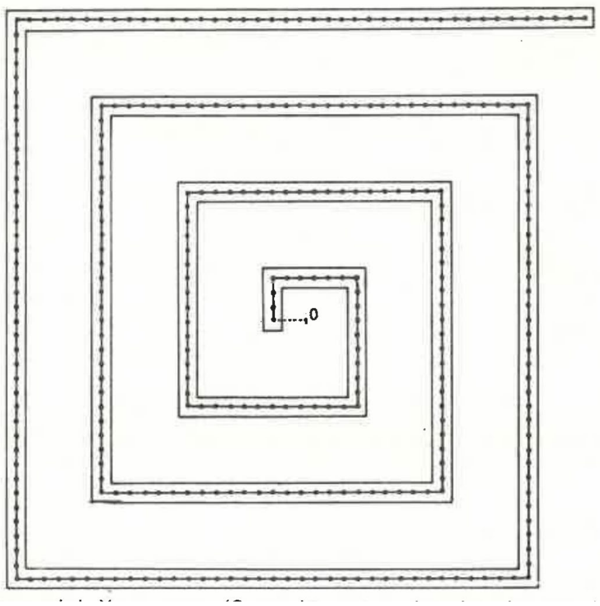

**Figure 2.** Jackson's square spiral. Young tsetse (*G. morsitans*) were released at the 'centre' (marked 0) and recaptured oh the spiral fly round. (The distance between dots corresponds to 100 yds; the perpendicular distance from the origin to the outer arm of the flyround was ±2100 yds).

In the second, so called 'large square' experiment, he used a 4 x 4 mile square of tsetse infested bush split into four sub-squares as shown in Fig.1. With this design, and by marking flies distinctively so that the day and sub-square of their release was known, it was possible (theoretically) to separate losses due to death and emigration respectively. Similarly, it was possible to separate between the effects of birth and immigration.

*Distance travelled from a point source; square spiral data*

Jackson (1946) noted the distance from the origin of the point of recapture of each fly. If one regards the recapture area as approximately a circle of radius $d = 2100$ yds then the data can be fitted to (9) in the text. (For this and all subsequent non-linear regressions the derivative-free programme PAR of the BMDP statistical package (Dixon and Brown, 1977) was used).

When the diffusion coefficient was assumed to be a constant ($k$) (so that $g = kt$) the model accounted for 79% of the variance (Fig. 3), and the rather scattered nature of the small number of data points available (df = 7 for the fitted model) suggested that more complex forms of $g$ would not remove significantly more of the variance. In support of this, when a model was fitted where $g = at + bt^2 + ct^3$ ($a$, $b$, $c$ constants), $b$ and $c$ took values of the order of $10^{-6}$ which were not significantly different from zero. Nor was there any significant decrease in the residual sum of squares with the more complex model ($P > 0.05$, $F$ test).

The value of $k$ obtained by regression when inserted into (7) gives $E(R) = 175$ yds when $t = 1$ day. That.is to say the modal predicts that a male *G. morsitans* will have moved, on average, 175 yds from its starting point after one day's dispersal. This is close to the value of about 162 yds (Rogers, 1977; Hargrove, 1981) obtained via the discrete random walk approach. On inserting the value of $k$, found by regression, into (3) the expected probability of a surviving fly being within the limits of the spiral is found, for any time $t$ after the start of the experiment. This probability will clearly decline monotonically with time after release, as will the probability of a fly surviving until time $t$, and therefore the product of these two probabilities. Thus, if the spiral was sampled fairly uniformly throughout (as seems reasonable (Hargrove, 1981)) the number of recaptures should also decay monotonically with time after release. In fact, however; the peak in recaptures occurred 2-3 weeks after release, presumably because the probability of capture changed with the age of the fly (Hargrove, 1981). (Recall that in this experiment all flies were less than 48 h old at release).

It is not possible to find explicitly the function describing the age-related change in capture probability, because it is inextricably confounded with the probability of survival. However, the product of the two probabilities should be proportional to the recapture rate divided by the probability ($E(R_d^0)$) that the fly is still within the spiral area being sampled. When Jackson's (1946) data were treated in this way (Fig. 4) they were well fitted by a function of the form

$$N = k_1\, t \exp(-k_2\, t) \qquad (20)$$

where $N$ was the total catch of (male) flies on day t. The change in catches could be viewed, then, as the product of a linear increase with age in the flies' probability of capture on a man fly round, with an exponentially decreasing probability of survival (*i.e.*, a constant death rate), Biological considerations indicate that this attractively simple model may not be entirely adequate (Hargrove, 1981) – but the data available will not support a more complex form,

For Jackson's (1946) data, when the sampling region was regarded as a square of side 3750 yds, and the value of $k$ already estimated was inserted into (19), the calculated value of $P_s(t)$ was nearly identical to the value $E(R_d^0)$calculated above.

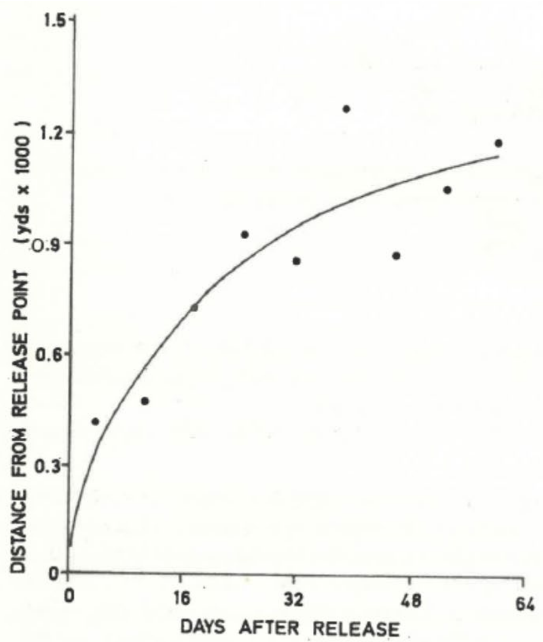


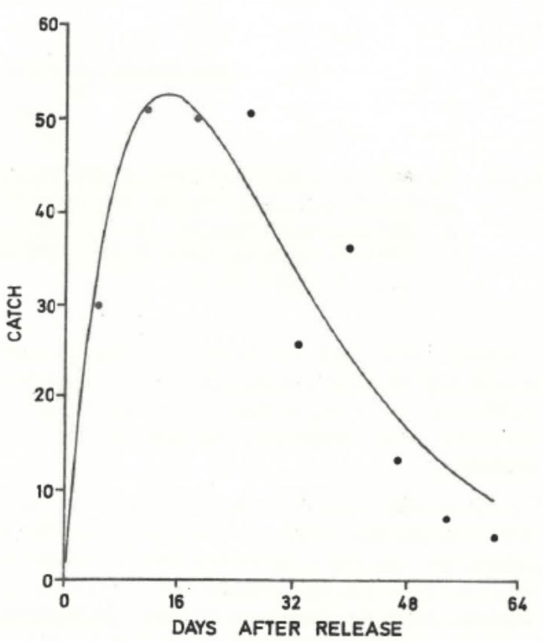


**Figure 3**. The apparent mean distance moved by male *G. morsitans* against time after release (at age ± 1 day). The line fitted is drawn according to Equation 9 with $R$ = 2.1 yds x $10^3$, $g(t) = kt$, and $k$ = 0.0195 (yds x $10^3)^2$/day. The asymptotic standard error for $k$ was 0.0028. Data from Jackson (1946).

**Figure 4**. Weekly catches of male *G. morsitans* from a rectangular spiral flyround; each catch was divided by the probability that a live fly of that age was still within the sampling area. The line fitted by non-linear regression was of the form $R = k_1 t \exp(-k_2 t)$, $k_1 = 10.08 \pm 1.37$, $k_2 = 7.03 \times 10^{-2} \pm 5.68 \times 10^{-3}$. (Units are days$^{-1}$ in each case).

*Diffusion out of a square: large square data*

The results obtained above indicate that it is sufficient to consider the diffusion coefficient for *G. m. morsitans* to be independent of the age of the fly. For the flies (of unknown age in this case) released in Jackson's l1941) large square experiment, the probability of a fly being in the square at time $t$ after release is given by the approximation (14), with $g = kt$ when the diffusion coefficient is a constant. When.one attempts to fit this function to Jackson's (1941) data it is obvious, though, that the shape of the function is quite inappropriate (Fig: 5). There is a discontinuity in the observed function, at the end of week 1 after release, which cannot be explained on the basis of the model chosen here.

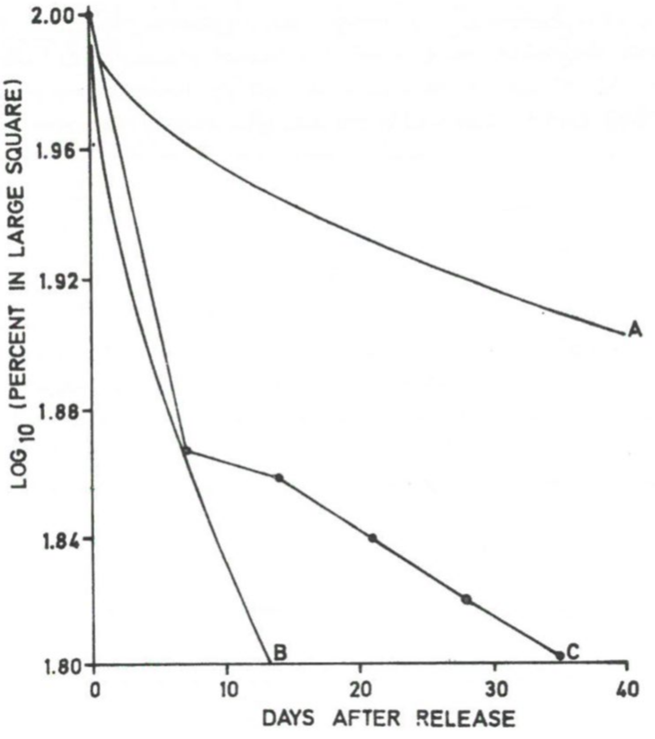


**Figure 5.** Theoretical and observed fraction of surviving population of male *G. morsitans* remaining within a square of side 4 miles, plotted against time after release from random points within the square.

A. Diffusion model (Equation 14), $k = 0.02$ (yds x $10^3)^2$.
B. Diffusion model (Equation 14), k = 0.20 (yds x $10^3)^2$.
C. Jackson's (1941) results.

Jackson (1941) noted this problem, and the fact that it could not have arisen as the result of a random walk process with constant step length. Various suggestions have been made over the years to explain away the discontinuity, or to give a reason for the departure of the results from those expected on the basis of random movement. Some explanations are probably the result of faulty logic (Hargrove, 1981); none seems wholly satisfactory.

Hargrove (1981) suggested that the observed loss from the square arose as the result of a complex interaction of a strong sampling bias (known to exist with the man fly-round system of sampling) with an increase in the rate of fly dispersal with age. Using *ad hoc* functions which incorporated this idea, he was able to obtain a good fit to the data shown in Fig. 5, This was achieved using simulation, however, and the correspondence at best only indicated the processes which may have been involved. The regression analysis of Jackson's (1946) 'square spiral' data in the previous section, produced no evidence in favour of the idea of a diffusion rate which increased with age, but the data available are too scanty to convince us that such a change could not exist. The 'large square' data, however, are actually consistent with the idea that the rate of diffusion slowed down with time after release, which would certainly not be evidence in favour of an increase of dispersal rate with age. On the other hand, because the flies were of unknown (and various) ages, and because of the complex changes in probability of capture with age (Hargrove, 1981), one should interpret the results with caution. A possible alternative explanation is that the early rapid dispersal is some manner of 'escape reaction', although Jackson (1941) argued that this was not the case since he was able to produce data indicating that the dispersal rate remained abnormally high for several days alter release. Furthermore, since flies were caught at random in the square, and released at their point of capture, there was no artificial increase in population density at any release point which could have been responsible for aberrant dispersal rates after release (Taylor, 1980).

Finally there is the possibility that the simple diffusion equation chosen here is, indeed, too much of a simplification, and that we need to consider more complex models. Taylor (1980) claims that data for the dispersal of *G. palpalis* and *G. tachinoides* conform better to a generalised gamma distribution than to the normal. Consideration of such models is beyond the scope of the present work, but we note that the more complex models involve an increased number of parameters and it is doubtful if the data considered here could provide an acceptable basis for a test of the suitability of such models.

#### ACKNOWLEDGEMENTS

This investigation received financial support from the UNDP/World Bank/WHO Special Programme for Research and Training in Tropical Diseases. K. Lange received support by way of an NIH Research Career Development Award (KO4 - HD00307) and from the Department of Biomathematics, University of California at Los Angeles. We thank John Van Sickle for helpful comments on the manuscript, Agnes Phiri for typing the original draft, and the Tsetse Control mapping office for preparing the figures. The work is published with the permission of the Director of Veterinary Services, Ministry of Agriculture, Zimbabwe.

#### REFERENCES

**Bursell, E.** (1970) Dispersal and concentration of *Glossina*. *in* The African Trypanosomiases (Ed. H. W. Mulligan). *pp* 382-394. London, George Allen and Unwin.

**Crumpacker, D. W. & Williams, J. S.** (1973) Density, dispersion and population structure in *Drosophila pseudoobsura*. *Ecological Monographs* **43**, 499-538.

**Dixon, W. J. & Brown, M. B.** (Ed.) (1977) Biomedical Computer Programmes. P-Series. Los Angeles, University of California Press.

**Dobzhansky, Th. & Wright, S.** (1943) Genetics of natural populations. X. Dispersion rates in *Drosophila pseudoobscura*. *Genetics*, **28**, 304-340.

**Hargrove**, **J.W.** (1981) Tsetse dispersal reconsidered. *Journal of Animal Ecology*, **50**, 351-373.

**Jackson, C. H. N.** (1941) The analysis of a tsetse fly population. *Annals of Eugenics*, **10**, 332-369.

**Jackson, C. H. N.** (1946) An artificially isolated generation of tsetse flies (Diptera). *Bulletin of Entomological Research*, **37**, 291-299.

**Kannan, D.** (1979) An introduction to stochastic processes. New York, Elsevier North Holland.

**Kareiva, P.M. (**1983) Local movement in herbivorous insects: applying a passive diffusion model to mark-recapture field experiments. *Oecologica*, **57**, 322-327.

**Rogers, D.** (1977) Study of a natural population of *Glossina fuscipes fuscipes* Newstead and a model of fly movement. *Journal of Animal Ecology*, **46**, 309-330.

**Skellam, J. G.** (1951) Random dispersal in theoretical populations. *Biometrika*, **38**, 196-218.

**Taylor, R. A. J.** (1980) A family of regression equations describing the density distributions of dispersing organisms. *Nature* **286**, 53-55.
